# ALDH Activity in Monocytes is Associated with Subclinical Coronary Atherosclerosis in Treated People with HIV-1

**DOI:** 10.64898/2026.08.20.746037

**Authors:** Jonathan Dias, Mohamed El-Far, Kevin Boczar, Abdelali Filali-Mouhim, Mehdi Benlarbi, Etiene Moreira Gabriel, Julie Moreaux, Soumia Khalfi, Tomas Raul Wiche Salinas, Marc Messier-Peet, Stéphane Isnard, Jean-Pierre Routy, Nicolas Chomont, Andrés Finzi, Carl Chantrand-Lefebvre, Cécile Tremblay, Madeleine Durand, Petronela Ancuta, the Canadian HIV and Aging Cohort Study (CHACS)

## Abstract

People living with HIV-1 (PWH) receiving antiretroviral therapy (ART) exhibit an increased cardiovascular disease (CVD) risk. Coronary atherosclerotic plaque formation is linked to chronic immune activation, a process fueled in PWH by additional mechanisms likely including residual HIV-1 production and microbial translocation from the gut during ART. Our previous studies demonstrated that aldehyde dehydrogenase (ALDH), an enzyme converting vitamin A into retinoic acid (RA), is upregulated in myeloid cells upon exposure to viral/bacterial/fungal products and that RA promotes HIV-1 production. To explore the link between ALDH/RA pathway and CVD risk, we used PBMC/plasma from ART-treated PWH (PWH^+^ART; n=51) and people without HIV-1 (Pw/oH; n=63) from the Canadian HIV/Aging Cohort Study, with/without subclinical coronary atherosclerosis, measured by coronary computed tomography angiography. ALDH expression was analyzed by flow cytometry on monocyte subsets, dendritic cells, and CD4^+^ T-cells. Unsupervised t-SNE-guided FlowSOM analysis associated top ALDH activity with a monocyte phenotype. The frequency of ALDH^+^ monocytes and plasma levels of RA and retinol binding protein 4 were increased in PWH^+^ART *versus* Pw/oH, with the highest RA levels coinciding with detectable plasma levels of soluble HIV-1 gp120. Multivariate regression models linked ALDH activity in monocytes to subclinical coronary atherosclerosis (*i.e.,* total plaque volume, low attenuated plaque volume, coronary artery calcification score) presence/burden in PWH^+^ART, independently of Framingham risk score. Finally, ALDH^+^ monocyte frequency and RA levels positively correlated with pericoronary fat attenuation index, an emerging CVD predictor. Thus, ALDH/RA pathway may represent a new marker of metabolic/immune dysfunction contributing to CVD risk in ART-treated PWH.

**KEY POINTS:**

- ALDH activity in blood monocytes is increased in people with HIV receiving antiretroviral therapy (ART) compared to people without HIV
- The frequency of ALDH+ monocytes is positively correlated with the atherosclerotic plaque burden during ART-treated HIV infection

**Graphical abstract:** 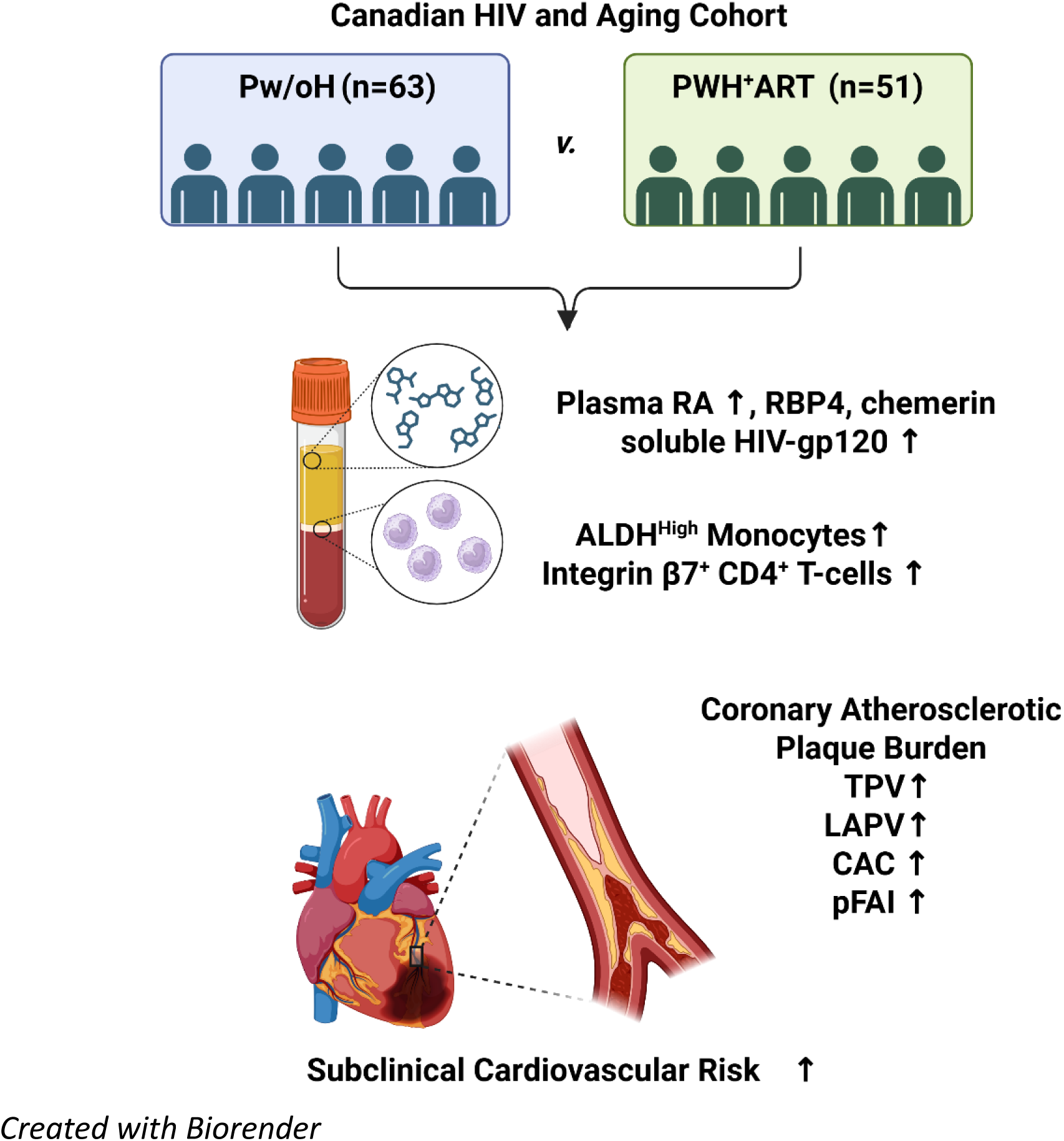

## Introduction

Antiretroviral therapy (ART) transformed human immunodeficiency type 1 (HIV-1) epidemic into a manageable chronic infection^1–5^. However, despite durable viral suppression, ART is not curative^1,3,4,6^. Integrated HIV-DNA persist during ART, viral rebound occurs following treatment interruption^3,4,7^, and people with HIV-1 (PWH) on ART experience an accelerated onset of age-associated comorbidities, particularly cardiovascular disease (CVD)^8–13^. Mechanisms linking HIV-1 reservoir persistence to the increased CVD risk in PWH on ART remain to be elucidated^14,15^.

Atherosclerosis develops progressively through chronic vascular inflammation, infiltration of immune cells into the arterial walls, and lipid accumulation within macrophages, leading to foam cell formation, myocardial infarction and stroke^16–19^. Single-cell RNA-sequencing identified immune cells infiltrating atherosclerotic lesions as inflammatory macrophages, foam cells, dendritic cells (DC), activated T-cells, and regulatory T-cells^20–22^. Other studies documented the deleterious contribution of macrophages and DC to atherosclerotic plaque formation^23–25^ and the protective features of Tregs^26,27^. Although traditional CVD risk factors are incorporated into the Framingham Risk Score (FRS), which estimates the 10-year cardiovascular risk by integrating age, sex, hyperlipidemia, high blood pressure, diabetes, and smoking^28^, FRS underestimates CVD risk in PWH on ART^29–31^.

Several HIV-specific factors contribute to CVD pathogenesis^11,32,33^. Long-term exposure to specific ART regimens has been associated with metabolic abnormalities and CVD risk^10,11,34^, although newer antiretroviral (ARV) drugs, particularly integrase strand transfer inhibitors (INSTI), demonstrated improved metabolic safety^35^. Increasing evidence suggests that HIV-1 persistence *per se* contributes to cardiovascular pathology^36^. Current ARVs do not directly target HIV-1 transcription/translation within viral reservoirs^37–41^; thus, residual production of HIV-1 RNAs/proteins/virions may sustain inflammatory pathways associated with vascular dysfunction during ART. Consistently, higher levels of cell-associated HIV-DNA/RNA have been associated with progression of coronary atherosclerosis in PWH on ART^42^, while residual viral transcription has been linked to monocyte dysfunction and vascular inflammation^35,43^. HIV-1 proteins, *e.g.,* gp120 and Nef, promote endothelial dysfunction, immune activation, and accelerated plaque development^44,45^. Studies by our group demonstrated that ART-treated PWH with subclinical coronary atherosclerosis display the highest levels of HIV-DNA in CD4^+^ T-cells^46^ and plasma soluble HIV-gp120 (sgp120)^47^. Furthermore, we identified enriched HIV-DNA levels in CD4^+^ T-cells with heart-homing phenotype^48^, pointing to the potential recruitment of HIV-1 reservoir cells in atherosclerotic plaques. Most recent studies revealed the contribution of HIV-infected myeloid cells to heart inflammation^19^.

A hallmark of HIV-1 pathogenesis is chronic immune activation linked to the incomplete restoration of mucosal barrier functions^49,50^. Microbial translocation from the gut perpetuates intestinal dysfunction and immune dysregulation during ART^5,49,51^. We previously demonstrated that bacterial products promote HIV-1 replication/viral outgrowth *via* mechanisms dependent on aldehyde dehydrogenase (ALDH) activity in myeloid cells^52^. The ALDH superfamily includes several isoforms (*e.g.,* ALDH1A1-3) involved in the conversion of retinol/vitamin A into retinoic acid (RA)^53^. Intestinal CD103^+^ DC express ALDH1A2 and represent a key source of RA in the intestinal environment^54,55^. We identified CD16^+^ monocytes as precursors of CD103^+^ DCs expressing ALDH activity *in vitro* and promoting HIV-1 replication/outgrowth in CD4^+^ T-cells *via* ALDH/RA-dependent mechanisms^52,56^. ALDH activity is induced by exposure to bacterial/fungal pathogens^51,52^, thus suggesting a link between microbial translocation and ALDH/RA pathways.

RA sustains mucosal immunity and contribute to HIV-1 pathogenesis^51^. RA signals through retinoic acid receptor α (RARα), a nuclear receptor that forms a heterodimer with RXR and binds onto RA-responsive elements (RARE) in the promoter regions of target genes^57^. We previously demonstrated that RA enhance HIV-1 replication and viral outgrowth in CD4^+^ T-cells and macrophages through mechanisms dependent on mTOR and SAMHD1^58,59^. Furthermore, the HIV-1 promoter contains RARE, and the recruitment of RARα:RXR heterodimers directly regulate HIV-1 transcription^59,60^. Beyond HIV-1 infection, the ALDH/RA pathways are documented to be involved in CVD pathogenesis^53,61–63^. Moreover, retinol-binding protein 4 (RBP4), the principal circulating carrier of retinol, regulates retinoid bioavailability to immune and vascular cells, has been associated with increased CVD risk^64,65^, and was recently demonstrated to reactivate latent HIV-1^66^. Furthermore, the RA downstream target chemerin, an adipokine involved in monocyte/macrophage migration, has also been associated with atherosclerosis severity and adverse cardiovascular outcomes^67,68^. Whether ALDH/RA pathway exacerbates the CVD risk in PWH on ART remains unknown.

In this study, we investigated the relationship between ALDH/RA pathway and subclinical coronary atherosclerosis in ART-treated PWH enrolled in the Canadian HIV and Aging Cohort Study (CHACS). Our findings identify increased ALDH activity in circulating monocytes as a novel immunometabolic feature associated with subclinical coronary atherosclerosis in PWH on ART and support the contribution of ALDH/RA-associated pathways to the CVD risk during ART-treated HIV-1 infection.

## Material and Methods

### Study Participants

The cardiovascular imaging sub-study of our CHACS cohort has been described previously^69–74^. Participants were free of prior cardiovascular events, had a 10-year FRS ranging between 5% and 20%, no known allergy to contrast medium and no renal failure. Participants underwent coronary computed tomography angiography (CCTA), with/without contrast, for the visualisation of coronary atherosclerotic plaque and were followed for clinical/laboratory investigations (Table 1-2). A total of n=114 CHACS participants were available for this study (n=63 Pw/oH; n=51 PWH^+^ART), with available PBMC samples (n=82) or plasma samples (n=79). Matched PBMC/plasma samples were available from n=34 PWH^+^ART and n=13 Pw/oH. PWH received ARVs: NRTI, PI, non-nucleoside reverse transcriptase inhibitors (NNRTI), and/or INSTI (<u>Supplemental File 1</u>). Collected blood was used to isolate PBMC and plasma, which were stored until use cryopreserved in media containing 10% DMSO in fetal bovine serum (FBS) in liquid nitrogen and at −80 °C, respectively.

**Table 1:**
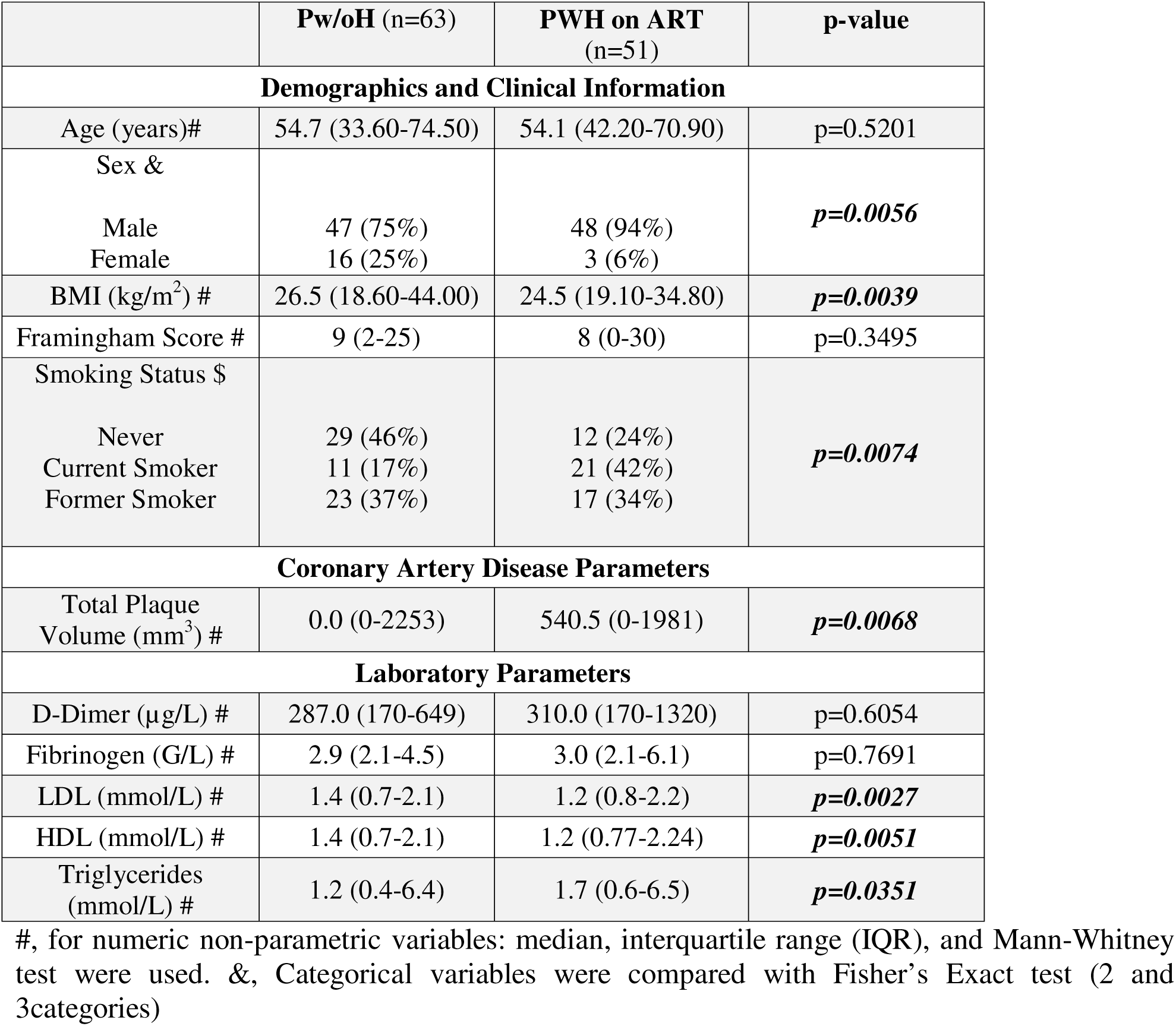
Description of Study Participants.

### Coronary computed tomography angiography

Total plaque volume (TPV) and low attenuated plaque volume (LAPV) were measured through contrast CCTA, while coronary artery calcification (CAC) score was measured through non-contrast CCTA, as previously described^69–74^. Pericoronary fat attenuation index (pFAI), a CCTA-derived marker of coronary vascular inflammation based on the attenuation characteristics of adipose tissue surrounding the coronary arteries, was quantified as previously described^75,76^.

#### Reagents

The list of reagents used is detailed in Supplemental Table 1.

### ALDEFLUOR ALDH Assay and Flow Cytometry Analysis

ALDH activity in PBMCs was measured using the ALDEFLUOR^TM^ assay (Supplemental Table 1), according to the manufacturer’s protocol (StemCell), as we previously reported^52^. Briefly, 1×10^6^ PBMCs were resuspended in 1 mL ALDEFLUOR assay buffer containing 5 µL of activated BODIPY aminoacetaldehyde (BAAA). Half of the cell suspension was immediately transferred to a tube containing the ALDH inhibitor (N,N-diethylaminobenzaldehyde; DEAB). PBMCs were incubated at 37°C for 45 minutes and subsequently washed with ALDEFLUOR assay buffer to prevent the efflux of BAAA. Cells were further incubated with antibodies against the CD3, CD4, CD16, CD14, CD1c, and HLA-DR (Supplemental Table 1). Dead cells were excluded using the Live/Dead Fixable Aqua Vivid Dead Cell Stain Kit (Thermo Fisher). Positive gates for ALDH activity were defined relative to the DEAB-treated negative control, as we reported^52^. Positive gates for Abs were defined using the fluorescence minus one (FMO) controls, as reported^77^. Samples were acquired using an LSRIIA flow cytometer with BD FACSDiva software (BD Bioscience). ALDH expression was analyzed using Flowjo software (Tree Star). In parallel, t-Distributed Stochastic Neighbour Embedding (t-SNE) and Self-Organizing Map (FlowSOM) analyses were performed on compensated live single cells using CellEngine (CellCarta) with the following parameters: 50% subsampling, channels including CD3, CD4, CD1c, HLA-DR, CD14, CD16; t-SNE perplexity=100; and number of nearest neighbours (k)=300. All remaining parameters were maintained at default settings. In addition, surface staining with ITGB7 Abs (Supplemental Table 1) was performed on CD4^+^ T-cells independently of ALDH activity staining.

### ELISA

Commercially available ELISA kits were used to quantify plasma concentrations of RA (MyBioSource), RBP4 (R&D Systems), and Chemerin (R&D Systems) (Supplemental Table 1).

### Soluble HIV-1 gp120 measurement

Plasma levels of sgp120 were quantified using a highly sensitive sandwich ELISA developed for the detection of circulating sgp120 in the plasma of PWH, as we described^47^.

### Statistical analysis

Statistical analyses were performed using GraphPad Prism version 11.0.1. Normal distribution was assessed using the Shapiro-Wilk test and prompted the use of non-parametric tests. Continuous variables were expressed as median and interquartile range (IQR). Comparisons between two unmatched groups were performed using the Mann-Whitney U test, whereas comparisons between two matched groups were performed using the Wilcoxon matched-pairs signed-rank test. Comparisons involving more than two unmatched groups were performed using Kruskal-Wallis tests followed by uncorrected Dunn’s multiple comparisons test. Comparisons involving more than two matched groups were performed using Friedman tests followed by uncorrected Dunn’s multiple comparisons test. Fisher’s exact tests were used for categorical comparisons involving two or more groups. Correlations were assessed using Spearman’s rank correlation coefficient. P-values <0.05 were considered statistically significant.

Zero-inflated gamma multivariate regression model as implemented in the glmmTMB package in R software (R Core Team 2019; https://www.R-project.org)^78^ was used to evaluate the predictive value of PBMC and plasma markers relative to cardiovascular imaging outcomes. Two complementary models were performed: the first identified predictors associated with plaque absence/presence, with TPV, LAPV and CAC used as categorical variables (<u>Supplemental Tables 2/4</u>), whereas the second identified predictors associated with plaque burden among study participants exhibiting detectable plaque, with TPV, LAPV and CAC used as continuous variables (<u>Supplemental Tables 3/5</u>). Crude models were adjusted for FRS (<u>Model 1</u>) and HIV-related parameters (<u>Model 2</u>; *i.e.,* duration of ART, duration of HIV-1 infection, nadir CD4 count, and CD4/CD8 ratio). To quantify the influence of confounding variables, the percentage change in odds ratio (%ΔOR) was calculated before and after covariate adjustment. Variations <10% were considered negligible, whereas shifts >30% were considered indicative of substantial confounding bias^78^. Multiple comparisons were corrected using the Benjamini– Hochberg procedure with a false discovery rate threshold defined by adjusted p-values, as described^72,73^.

## Results

### Clinical and laboratory profiles of the CHACS participants

Participants from the CHACS cohort were classified based on the HIV-1 status and the presence (TPV^+^; TPV>0 mm^3^) or the absence (TPV^−^; TPV=0 mm^3^) of subclinical coronary atherosclerosis, as we previously described^72,73^. Pw/oH and PWH^+^ART were similar in age, FRS, D-dimer, and fibrinogen, but differed in male/female distribution (*p=0.0056*), smoking prevalence (*p=0.0074*), body mass index (BMI) (p*=0.0039*), TPV (*p=0.0068*), low-density lipoprotein (LDL) (*p=0.0027*), high-density lipoprotein (HDL) (*p=0.0051*), and triglycerides (*p=0.0351*) (Table 1). Among PWH^+^ART participants with *versus* without subclinical coronary atherosclerosis only differed in fibrinogen levels (*p=0.0329*) and the ART duration (*p=0.0257*) (Table 2).

**Table 2:** Clinical Parameters of PWH on ART Study Participants with and without Subclinical Atherosclerosis.

|  | TPV<sup>-</sup> (n=19) | TPV<sup>+</sup> (n=29) | p-value |
| --- | --- | --- | --- |
| **Demographics and Clinical Information** |  |  |  |
| Age (years) # | 53.5 (42.2-62.1) | 54.1 (44.2-70.9) | p=0.2689 |
| Sex & |  |  | p>0.9999 |
| Male | 18 (95%) | 27 (93%) |  |
| Female | 1 (5%) | 2 (7%) |  |
| BMI (kg/m<sup>2</sup>) # | 25.7 (21-34.8) | 23.5 (19.1-34.5) | p=0.1346 |
| Framingham Score # | 7 (3-18) | 9 (0-30) | p=0.0873 |
| Smoking & |  |  | p=0.1127 |
| Never | 5 (28%) | 3 (11%) |  |
| Current Smoker | 5 (28%) | 16 (59%) |  |
| Former Smoker | 8 (44%) | 8 (30%) |  |
| **Laboratory Parameters** |  |  |  |
| D-Dimer (ug/L) # | 290.0 (170-1320) | 340 (170-720) | p=0.5623 |
| Fibrinogen (G/L) # | 2.9 (2.1-3.7) | 3.2 (2.1-6.1) | **p=0.0329** |
| LDL (mmol/L) # | 2.9 (1.9-4.6) | 2.4 (0.8-8.0) | p=0.1991 |
| HDL (mmol/L) # | 1.2 (0.8-2.0) | 1.2 (0.8-2.2) | p=0.8981 |
| Triglycerides (mmol/L) # | 1.7 (0.8-3.2) | 1.9 (0.6-6.5) | p=0.2099 |
| **Serology** |  |  |  |
| Co-infection CMV & |  |  | p=0.6376 |
| Yes | 15 (94%) | 21 (88%) |  |
| No | 1 (6%) | 3 (12%) |  |
| **HIV-1 Disease Parameters** |  |  |  |
| Duration of HIV-1 (years) # | 17.5 (3.8-29.4) | 19.2 (5.7-30.8) | p=0.3002 |
| Duration of ART (years) # | 12.7 (0.0-25.0) | 16.7 (3.3-24.8) | **p=0.0257** |
| Nadir CD4 (x10<sup>9</sup>/L) # | 200 (10.0-750.0) | 230 (10.0-576.0) | p=0.8657 |
| CD4 (%) # | 33.0 (13.0-57.0) | 32.0 (3.0-46.0) | p=0.5804 |

#, for numeric non-parametric variables: median, interquartile range (IQR), and Mann-Whitney test were used. &, Categorical variables were compared with Fisher's Exact test (2 and 3 categories).

### Enhanced ALDH activity in HLA-DR^+^CD14^+^ monocytes in PWH^+^ART versus Pw/oH

The ALDEFLUOR assay was used to identify PBMC subsets exhibiting ALDH activity in Pw/oH and PWH^+^ART CHACS participants (Figure 1A). Unsupervised FlowSOM followed by t-SNE analyses identified five major immune populations, with ALDH activity predominantly enriched in HLA-DR^+^CD14^+^ monocytes (FlowSOM.10.10), followed by HLA-DR^+^CD1c^+^ DC (FlowSOM.10.05), and minimal ALDH expression in CD3^+^CD4^+^ T-cells (FlowSOM.10.08), CD4^−^CD3^+^ T-cells (FlowSOM.10.03), and CD3^−^CD14^−^CD1c^−^CD16^+^ cells (FlowSOM.10.02) (Figure 1B-C). The frequency of FlowSOM.10.10/ALDH^+^HLA-DR^+^CD14^+^ was significantly increased in PWH^+^ART compared with Pw/oH (*p=0.0007)* (Figure 1D), identifying monocytes as the major ALDH^+^ population disproportionately expanded in PWH^+^ART.

**Figure 1.**
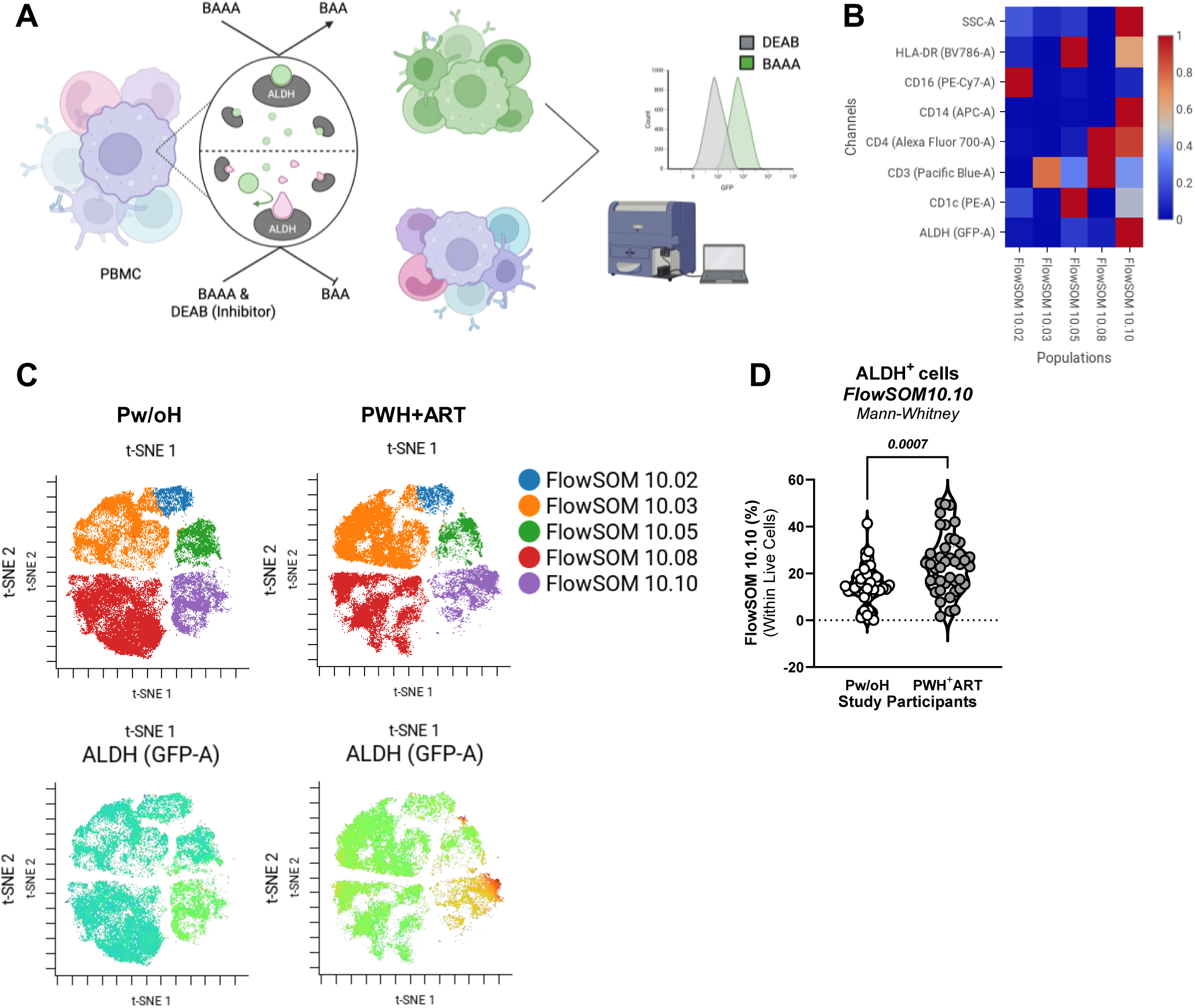
Characterization of ALDH activity in PBMCs of CHACS participants using the ALDEFLUOR assay and unsupervised flow cytometry analysis. **(A)** PBMCs from the CHACS cohort were analyzed for ALDH activity using the ALDEFLUOR™ assay. Briefly, cells were incubated with the ALDH substrate BAAA in the presence or absence of the selective ALDH inhibitor DEAB, which was used to define background fluorescence. In parallel, extracellular staining was performed using antibodies against CD3, CD4, CD1c, HLA-DR, CD14, and CD16, together with a Live/Dead viability dye, to phenotypically identify major immune cell populations by flow cytometry. **(B)** Unsupervised clustering of live PBMCs using FlowSOM generated a heat map summarizing marker expression profiles across identified immune cell clusters. **(C)** Representative t-SNE projections from one Pw/oH and one PWH^+^ART illustrating the distribution of PBMC populations according to FlowSOM clustering, with ALDH activity overlaid onto the clustering topology. **(D)** Quantification of FlowSOM cluster 10.10, corresponding to the ALDH-enriched HLA-DR^+^CD14^+^ monocyte population, comparing its relative frequency between Pw/oH (n=40) and PWH^+^ART (n=42). Statistical significance was determined using the Mann–Whitney test, with p-values indicated on the graph.

### Preferential contribution of classical monocytes to the pool of ALDH^+^ monocytes

Manual gating was further used to characterize the distribution of ALDH activity across SSC^high^ monocytes and DCs (Supplemental Figure 1A). PWH^+^ART *versus* Pw/oH exhibited increased frequencies of total monocytes and reduced frequencies of DC and CD4^+^ T-cells (*p<0.0001*) (Supplemental Figure 1B-D). Consistent with the FlowSOM analysis (Figure 1), PWH^+^ART *versus* Pw/oH exhibited increased frequencies of SSC^High^ALDH^+^ cells and ALDH MFI (Supplemental Figure 2A-C). Within the SSC^High^ cells, monocytes expressed higher ALDH activity compared to DCs (Figure 2A-B), with the frequency of ALDH^+^ monocytes and DCs being significantly increased in PWH^+^ART *versus* Pw/oH (*p=0.0010*) (Figure 2C). A tendency for increased ALDH^+^ frequency in DCs (*p=0.0789*) (Figure 2C) and no differences in ALDH MFI for monocytes and DCs (Figure 2D) were observed in PWH^+^ART *versus* Pw/oH. Finally, the pool of ALDH^+^ myeloid cells was mainly composed of monocytes *versus* DCs (Figure 2E).

**Figure 2.**
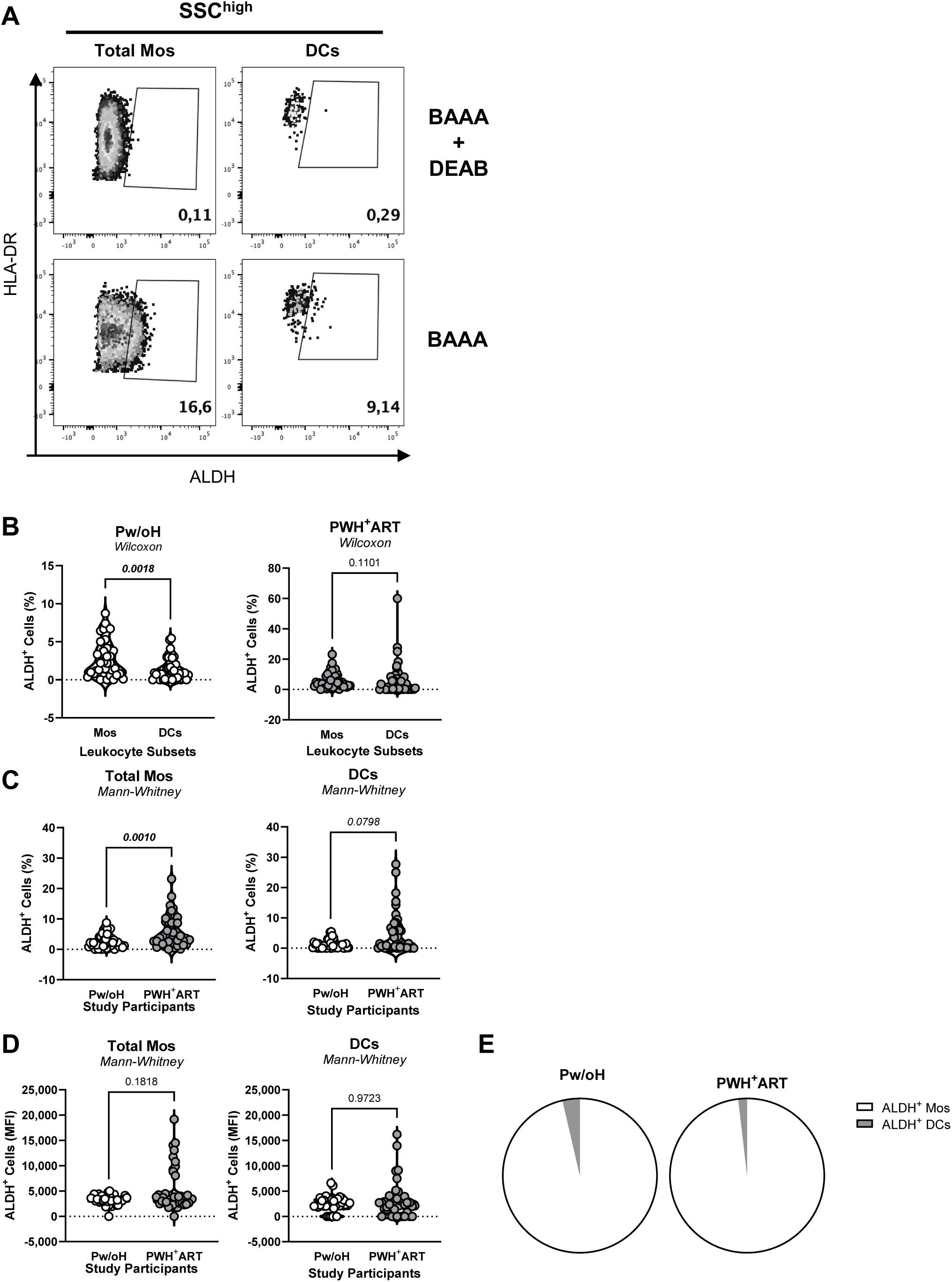
ALDH activity in monocytes and DC of CHACS participants. PBMCs from Pw/oH and PWH^+^ART were stained for surface markers and ALDH activity as described in Figure 1. ALDH activity in PBMC subsets was assessed by flow cytometry using manual gating performed in FlowJo, with total monocytes and DC identified within the SSC^High^ compartment. **(A)** Representative flow cytometry plots illustrating ALDH^+^ total monocytes and DC following BAAA staining relative to DEAB-treated controls. **(B)** Frequencies of ALDH^+^ total monocytes and DC for Pw/oH (left panel) and PWH^+^ART (right panel). **(C)** Comparison of ALDH^+^ total monocyte frequencies (left panel) and ALDH^+^ DC frequencies (right panel) between Pw/oH and PWH^+^ART. **(D)** MFI of ALDH activity within total monocytes (left panel) and DCs (right panel) comparing Pw/oH and PWH^+^ART. Two-group comparisons were performed using the Mann– Whitney test (panels C-D), whereas comparisons between monocytes and DCs within the same study group were performed using the Wilcoxon matched-pairs signed-rank test (panel B). P-values are indicated on the graphs.

Further, ALDH activity was analysed in monocyte subsets identified based on their differential CD14/CD16 expression (Supplemental Figure 3A). A significant increase was observed in the frequency of classical (CD14^+^CD16^−^; *p=0.0392*), and a decrease in intermediate (CD14^+^CD16^+^; *p=0.0128*) and non-classical monocytes (CD14^−^CD16^+^; *p<0.0001*) in PWH^+^ART *versus* Pw/oH, with classical monocytes representing the predominant subset (Supplemental Figure 3B). Intermediate and non-classical *versus* classical monocytes exhibited the highest ALDH activity (Figure 3A-B). However, classical monocytes were the most abundant and represented the predominant contributor to the ALDH^+^ pool in Pw/oH and PWH^+^ART (Figure 3C). PWH^+^ART *versus* Pw/oH exhibited increased frequencies of ALDH^+^ classical (*p=0.0128*), intermediate (*p=0.0051*), and non-classical monocytes (*p=0.0002*), without significant differences in ALDH MFI (Figure 3D-E). Thus, ALDH activity is increased in PWH on ART compared to Pw/oH, with classical monocytes contributing predominantly to the pool of ALDH^+^ cells.

**Figure 3.**
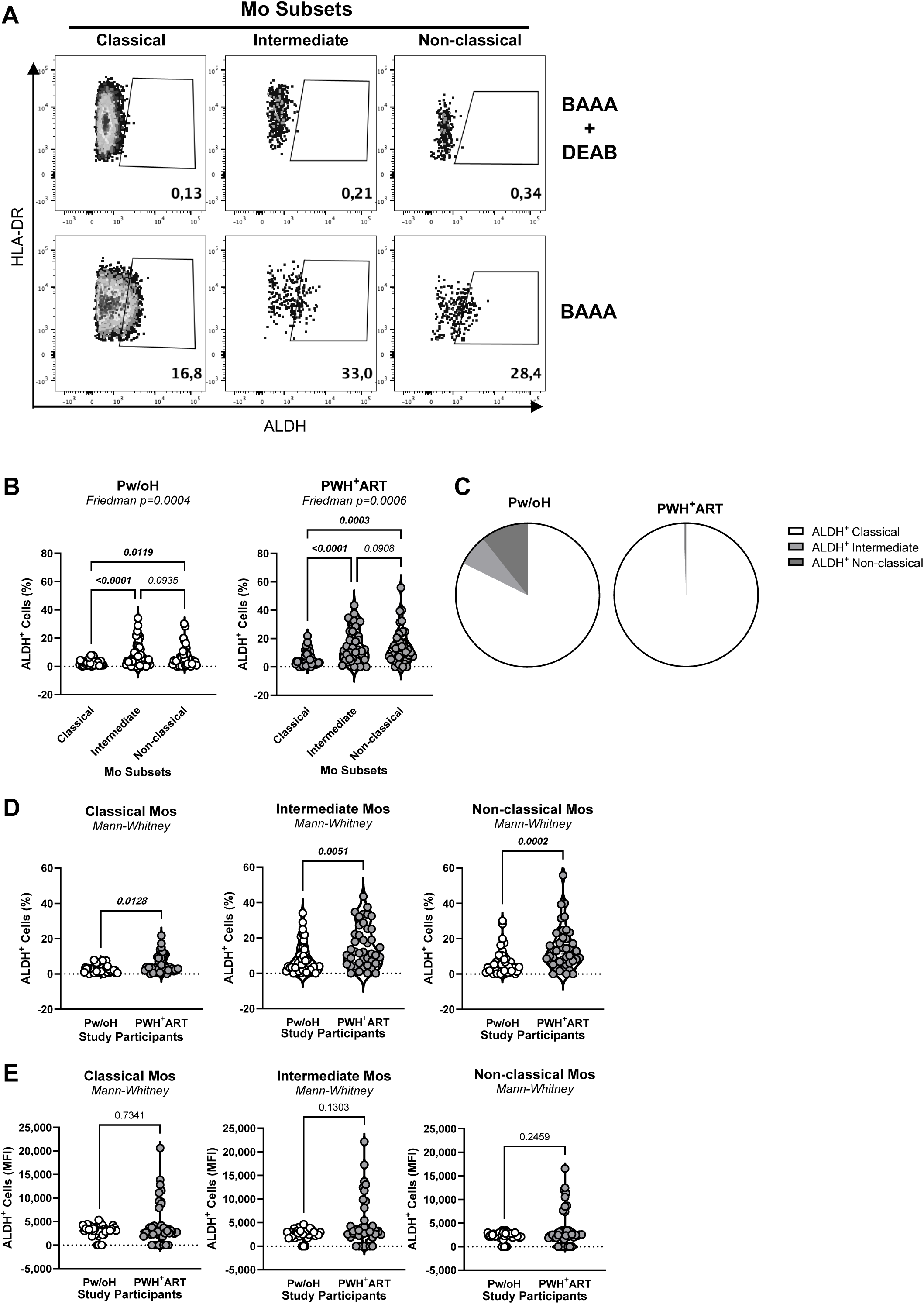
Differential ALDH activity across monocyte subsets of CHACS participants. PBMCs from Pw/oH and PWH^+^ART were stained for surface markers and ALDH activity as described in the Figure 1. Monocyte subsets were identified by manual gating according to CD14 and CD16 expression profiles, with classical monocytes defined as CD14^+^CD16^−^, intermediate monocytes as CD14^+^CD16^+^, and non-classical monocytes as CD14^−^CD16^+^. **(A)** Representative flow cytometry plots illustrating ALDH^+^ classical, intermediate, and non-classical monocyte subsets following BAAA staining relative to DEAB-treated controls. **(B)** Frequencies of ALDH^+^ classical, intermediate, and non-classical monocyte subsets within Pw/oH (left panel) and PWH^+^ART (right panel). **(C)** Pie chart representation illustrating the relative contribution of classical, intermediate, and non-classical monocyte subsets to the total ALDH^+^ monocyte compartment in Pw/oH and PWH^+^ART. **(D)** Comparison of ALDH^+^ classical (left panel), intermediate (middle panel), and non-classical (right panel) monocyte frequencies between Pw/oH and PWH^+^ART study participants. **(E)** MFI of ALDH activity within classical (left panel), intermediate (middle panel), and non-classical (right panel) monocyte subsets comparing Pw/oH and PWH^+^ART. Comparisons among monocyte subsets within the same study group were performed using Friedman tests followed by uncorrected Dunn’s multiple comparisons tests (panel B). Comparisons between Pw/oH and PWH^+^ART were performed using Mann– Whitney tests (panels D-E). p-values are indicated on the graphs.

### Increased ITG*β*7 expression in SSC^Low^ leukocytes and CD4^+^ T-cells of PWH^+^ART

To determine the impact of ALDH activity on the expression of RA-target genes, we investigated the expression of ITGB7, a RA-modulated gut-homing molecule^51,58^. In contrast to ALDH expression mainly observed on SSC^High^ myeloid cells (Supplemental Figure 2A), ITGB7 was mainly expressed on SSC^Low^ lymphoid cells (Figure 4A). PWH^+^ART *versus* Pw/oH exhibited increased frequencies of SSC^Low^ITGB7^+^ cells (*p=0.0025*), but similar ITGB7 MFI (Figure 4B). SSC^Low^ITGB7^+^ cell frequencies positively correlated with the frequencies of SSC^High^ALDH^+^ cell in Pw/oH (*r=0.3243; p=0.0412*) but not PWH^+^ART (Figure 4C). The frequency (*p=0.0275*) and MFI (*p=0.0292*) of ITGB7 expression on CD4^+^ T-cells (Figure 4D) were further analysed and demonstrated increase in PWH^+^ART *versus* Pw/oH (Figure 4E). Finally, the correlation between the frequencies of ITGB7^+^ CD4^+^ T-cells and SSC^High^ALDH^+^ cells did not reach statistical significance in PWH^+^ART, despite a tendency for Pw/oH (Figure 4F). Thus, ITGB7 expression is elevated in PWH^+^ART *versus* Pw/oH, with ITGB7 and ALDH activity in the blood compartment being positively correlated in Pw/oH but not PWH^+^ART.

**Figure 4.**
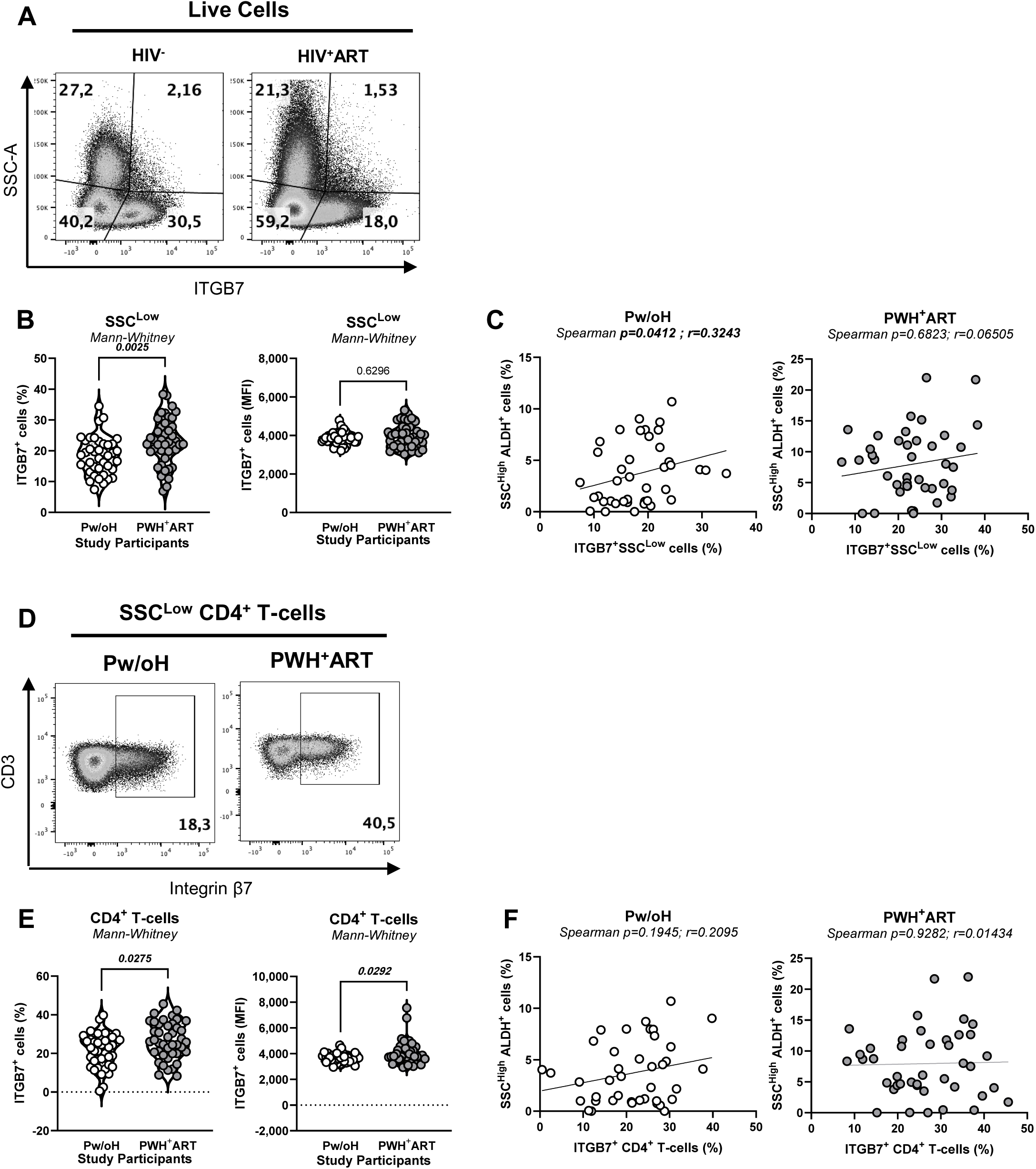
ITGB7 expression in leukocyte populations and CD4^+^ T-cells of CHACS participants. PBMCs from Pw/oH and PWH^+^ART were analyzed for ITGB7 expression by flow cytometry. **(A)** Representative flow cytometry plots illustrating ITGB7 expression across live leukocyte populations according to SSC properties in Pw/oH and PWH^+^ART study participants. **(B)** Frequency (left panel) and MFI (right panel) of ITGβ7^+^ cells within the SSC^Low^ leukocyte compartment comparing Pw/oH and PWH^+^ART. **(C)** Correlation analyses between the frequency of ALDH^+^ SSC^High^ leukocytes and the frequency of ITGB7^+^ SSC^Low^ leukocytes in Pw/oH (left panel) and PWH^+^ART (right panel). **(D)** Representative flow cytometry plots illustrating ITGβ7 expression within SSC^Low^ CD3^+^CD4^+^ T-cells in Pw/oH and PWH^+^ART. (E) Frequency (left panel) and MFI (right panel) of ITGB7^+^ CD4^+^ T-cells comparing Pw/oH and PWH^+^ART. **(F)** Correlation analyses between the frequency of ALDH^+^ SSC^High^ leukocytes and the frequency of ITGB7^+^ CD4^+^ T-cells in Pw/oH (left panel) and PWH^+^ART (right panel). Comparisons between Pw/oH and PWH^+^ART were performed using Mann–Whitney tests (panels B and E). Correlation analyses were performed using Spearman rank correlation tests (panels C and F). Correlation coefficients (r) and p-values are indicated on the graphs.

### Coronary artery atherosclerosis is associated with monocyte ALDH activity in PWH^+^ART

The subclinical coronary atherosclerotic plaque burden was measured in Pw/oH and PWH^+^ART, with TPV reflecting cumulative coronary plaques burden^71^, LAPV representing lipid-rich plaque components associated with increased cardiovascular risk^79^, and CAC serving as a marker of long-term cardiovascular risk^80^. Among participants with detectable coronary plaque, TPV (*p=0.0128*), LAPV (*p=0.0104*) and CAC values (*p=0.0380*) were significantly elevated in PWH^+^ART *versus* Pw/oH (Figure 5A). A zero-inflated gamma regression modeling were used to determine the relationship between plaque presence/burden and ALDH activity in myeloid cells and ITGB7 expression in lymphoid cells (Supplemental Table 2-5). After adjustment for FRS, ALDH activity within SSC^High^ myeloid cells was associated with plaque presence defined by TPV, LAPV, and CAC, although these associations were marginal after multiple-comparison adjustment (Supplemental Table 2). Total monocyte ALDH activity was similarly associated with TPV (*Adj. p=0.0465*) and LAPV (*Adj. p=0.0465*), with a trend for CAC (*p=0.0304; Adj. p=0.0608* (Supplemental Table 2). Similar but marginally significant associations were observed for classical and intermediate monocytes (Supplemental Table 2). Following adjustment for HIV-related parameters, only ALDH activity in intermediate monocyte retained nominal associations with TPV, LAPV, and CAC, which did not remain significant after multiple-comparison adjustment (Supplemental Table 2).

**Figure 5.**
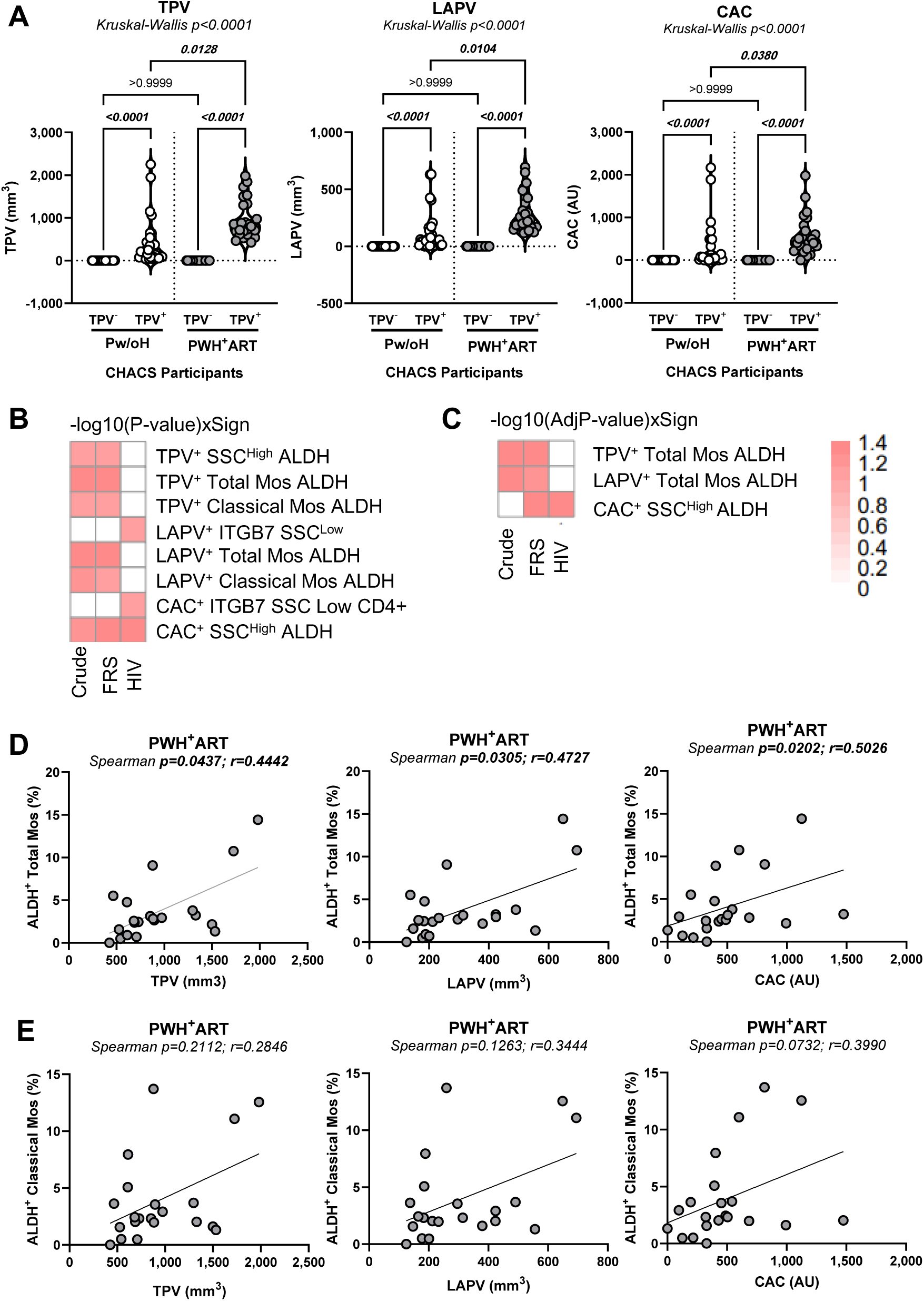
Association between ALDH activity and coronary atherosclerotic plaque burden in CHACS participants. **(A)** Pw/oH and PWH^+^ART were stratified according to the absence (TPV^−^) or presence (TPV^+^) of detectable coronary atherosclerotic plaques quantified by CCTA. TPV (left panel), LAPV (middle panel), and CAC (right panel) measurements are shown for Pw/oH and PWH^+^ART. **(B-C)** Heat maps summarizing significant associations identified by logistic regression analyses between the atherosclerotic plaque burden (TPV, LAPV, and CAC) and ALDH activity within immune cell populations. **(B)** Heatmaps depicting significant crude model associations represented as -log10(p-value)×sign, whereas **(C)** shows associations remaining significant after adjustment analyses represented as -log10(adjusted p-value)×sign. Models were adjusted for FRS or HIV-related clinical parameters (Supplemental Tables 2-3). **(D)** Correlation analyses between ALDH^+^ total monocyte frequencies and TPV (left panel), LAPV (middle panel), or CAC (right panel) in PWH^+^ART study participants. **(E)** Correlation analyses between ALDH+ classical monocyte frequencies and TPV (left panel), LAPV (middle panel), or CAC (right panel) in PWH^+^ART study participants. Statistical analyses were performed using Kruskal-Wallis tests followed by uncorrected Dunn’s multiple comparisons tests. P-values are indicated on the graphs. Correlations were assessed using Spearman rank correlation tests. Correlation coefficients (r) and p-values are indicated on the graphs.

Among participants with detectable plaque, FRS-adjusted analyses demonstrated that ALDH activity in SSC^High^ myeloid cells was associated with CAC burden (*Adj. p=0.0387*), whereas ALDH activity in total monocytes was associated with TPV burden (*Adj. p=0.0467*) and showed a marginal association with LAPV burden (*Adj. p=0.0517*) (Figure 5B-C; <u>Supplemental Table 3</u>). ALDH activity in classical monocytes showed nominal associations with TPV and LAPV burden, which did not remain significant after multiple comparison adjustment (Figure 5B-C; <u>Supplemental Table 3</u>). Following adjustment for HIV-related parameters, only the association between SSC^High^ ALDH^+^ cells and CAC burden remained nominally significant (*Adj. p=0.0502*), whereas associations with total and classical monocytes lost significance (Figure 5B-C; <u>Supplemental Table 3</u>). No significant associations were observed for intermediate or non-classical monocytes or ITGβ7 expression (Figure 5B-C; <u>Supplemental Table 3</u>). Consistent with these models, ALDH activity in total monocytes positively correlated with TPV (*r=0.4442; p=0.0437*), LAPV (*r=0.4727; p=0.0305*), and CAC (*r=0.5026; p=0.0202*) burden in PWH^+^ART (Figure 5D), whereas correlations did not reach statistical significance for ALDH^+^ classical monocytes (Figure 5E). Together, these results associate ALDH activity in monocytes with coronary plaque presence and burden in ART-treated PWH.

### Retinoid pathway dysregulation in PWH^+^ART

We next investigated whether systemic components of the retinoid pathway were altered in relationship with the HIV/ART and CVD status. We quantified plasma RA, as a read-out of ALDH activity^53^, RBP4 involved in retinol transport^81,82^ and HIV-1 latency reversal^66^, and chemerin, another RA-modulated gene and CVD marker^83,84^. RA levels were significantly increased in PWH^+^ART *versus* Pw/oH (*p=0.0305*) but did not correlate with TPV, LAPV, or CAC burden (Figure 6A). RBP4 tented to be elevated in PWH^+^ART and positively correlated with TPV (*r=0.5607; p=0.0066*) and LAPV (*r=0.5641; p=0.0062*), but not CAC burden (Figure 6B). Chemerin levels did not differ in PWH^+^ART *versus* Pw/oH, and were not correlated with plaque (Figure 6C). In multivariable analyses, none of these plasma markers independently predicted plaque presence/burden after adjustment for FRS or HIV-related parameters, with the exception of RBP4 that showed a tendency toward significance (Supplemental Tables 4-5). Thus, PWH^+^ART exhibited systemic alterations in retinoid-associated biomarkers, with RBP4 levels emerging as potential new correlates of coronary atherosclerosis burden.

**Figure 6.**
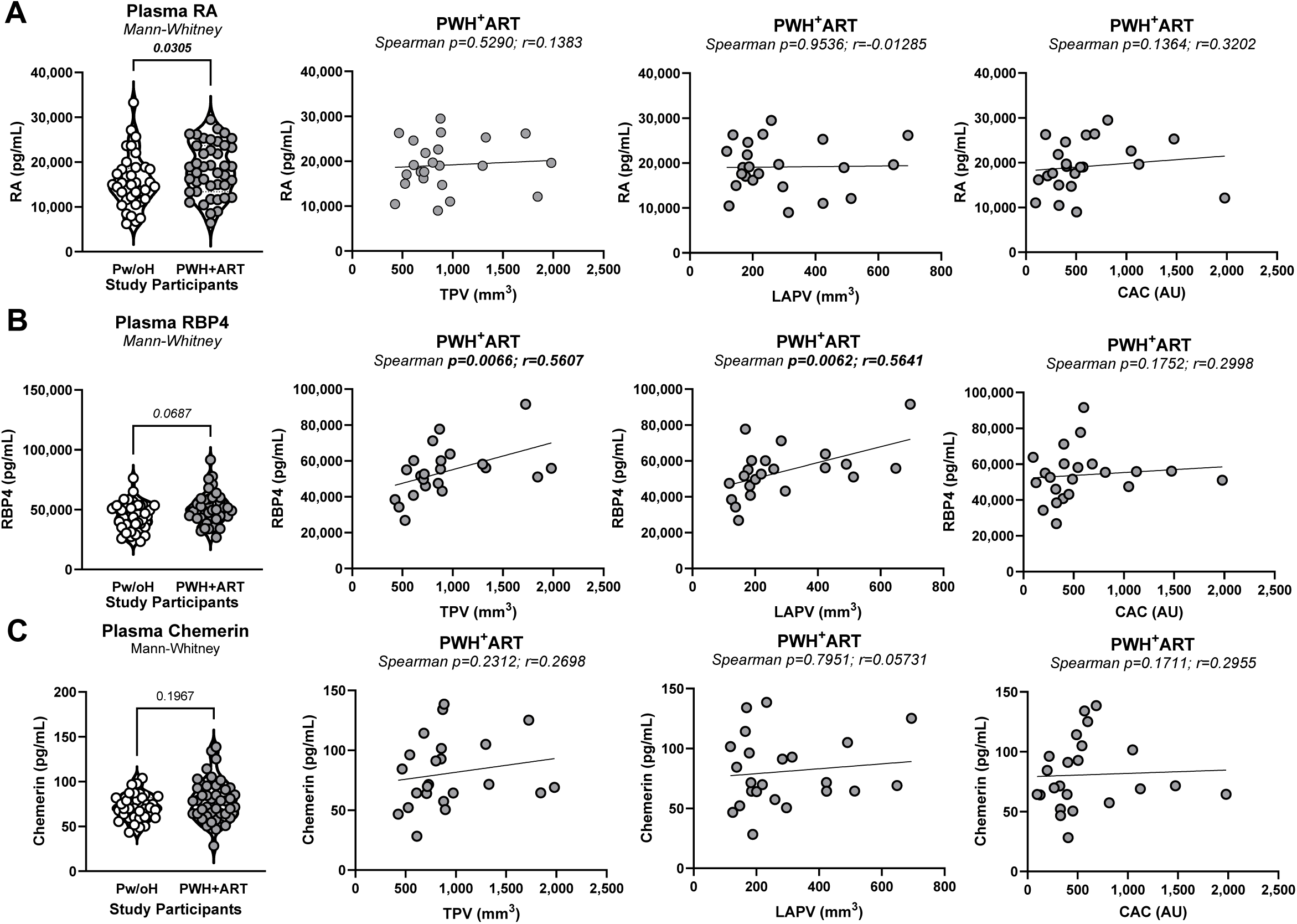
Plasma retinoid-associated biomarkers in CHACS participants. Plasma concentrations of (A) RA, (B) RBP4, and (C) chemerin were quantified and compared between Pw/oH and PWH^+^ART. Associations between plasma biomarker concentrations and TPV (left panel), LAPV (middle panel), and CAC (right panel) were subsequently evaluated in PWH^+^ART compared to Pw/oH. Comparisons between groups were performed using Mann–Whitney tests. Correlation analyses were performed using Spearman rank correlation tests. Correlation coefficients (r) and p-values are indicated on the graphs.

### Plasma soluble HIV-1 gp120 detection coincides with increased RA levels

PWH^+^ART were further stratified based on the presence/absence of detectable sgp120 and analyzed for differences in ALDH activity, and plasma levels of RA, RBP4 and chemerin. PWH^+^ART with detectable *versus* undetectable plasma sgp120 exhibited significantly higher plasma levels of RA (*p=0.0309*), but similar levels of ALDH, RBP4 and chemerin (Supplemental Figure 3A-D, <u>left panels</u>). Although correlations did not reach statistical significance (Supplemental Figure 3A-D, <u>right panels</u>), likely due to limited sample size, our findings point to a potential link between residual HIV-1 production and RA in ART-treated PWH.

### Associations between pFAI and retinoid pathway markers in PWH^+^ART

The pFAI, a recently proposed non-invasive imaging biomarker of coronary inflammation measured by CCTA^75,76^, was further examined for its associations with ALDH/retinoid pathway in CHACS participants (Figure 7). In Pw/oH, PFA values positively correlated with RA levels, but not with the frequency of ALDH^+^ total/classical monocytes, nor RBP4 and chemerin levels (Figure 7A-E, <u>left panels</u>). In PWH^+^ART, pFAI values only positively correlated with the frequency of ALDH^+^ classical monocytes and RA levels (Figure 7A-E, <u>right panels</u>). These findings link systemic RA levels and ALDH activity to pFAI-defined coronary inflammation in PWH on ART.

**Figure 7.**
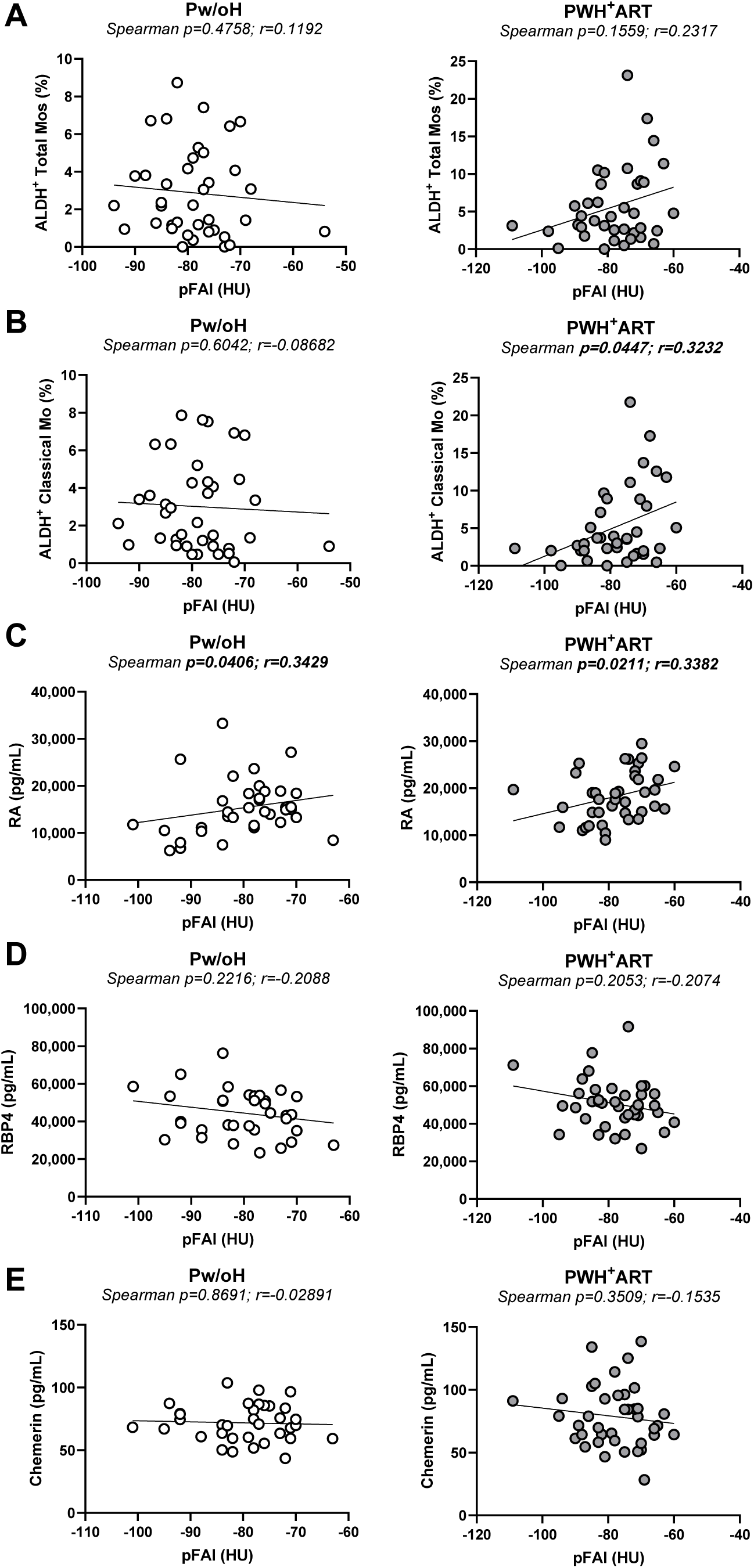
Associations between pFAI and retinoid pathway-associated biomarkers in CHACS participants. Correlation analyses between pFAI measurements and (A) ALDH^+^ total monocyte frequencies, (B) ALDH^+^ classical monocyte frequencies (C) plasma RA concentrations, (D) plasma RBP4 concentrations, and (E) plasma chemerin concentrations in Pw/oH (left panels) and PWH^+^ART (right panels) study participants. Correlations were assessed using Spearman rank correlation tests. Correlation coefficients (r) and p-values are indicated on the graphs.

### Impact of statin use on retinoid pathway markers and immune cell ALDH activity

Given previous evidence demonstrating interactions between statins and ALDH enzymes^85^, as well as mechanistic links between ALDH2 and HMG-CoA reductase activity^86,87^, we finally examined the impact of statin use on ALDH/RA pathway markers. No statistically significant differences were observed between Statin^+^ (n=12) *versus* Statin^−^ (n=30) groups in terms of frequency of ALDH+ monocytes, DCs, classical, intermediate and non-classical monocytes (Supplemental Figure 5A-C), ITGB7 expression on SSC^Low^ lymphoid cells (Supplemental Figure 5D), nor plasma RA and chemerin levels (Supplemental Figure 5E, <u>left/right panels</u>). Only RBP4 levels were significantly increased in Statin^+^ *versus* Statin^−^ PWH^+^ART (Supplemental Figure 5E, <u>middle panel</u>). Thus, statin treatment was not associated with a decrease in ALDH/RA pathway markers.

## Discussion

The present study identifies overt expression of the ALDH/RA pathway as a novel metabolic/immune dysfunction associated with subclinical coronary atherosclerosis in PWH on ART. These findings extend knowledge that emerged previously from our studies linking subclinical coronary atherosclerosis to alterations in circulating Th17-cells and monocytes^72^, gut-homing and tissue-resident T-cell populations^73^, and markers of HIV-1 persistence, including cell-associated HIV-DNA^46^ and plasma HIV-1 sgp120^47^.

Increasing evidence supports the existence of a gut-heart axis in CVD pathogenesis^88^, whereby impaired intestinal barrier integrity and microbial translocation contribute to systemic inflammation and vascular dysfunction^89–91^. The fact that bacteria/fungal compounds promote ALDH activity^52^ points to this metabolic feature as a new functional read-out for chronic exposure to microbial products and subsequent inflammation-associated CVD risk. In this study, ALDH activity was predominantly enriched in HLA-DR^+^CD14^+^ monocytes and was significantly increased across classical, intermediate, and non-classical monocytes, with classical monocytes predominantly contributing to the pool of ALDH^+^ myeloid cells in ART-treated PWH. Consistently, PWH on ART distinguished from Pw/oH by increased plasma RA and RBP4 concentrations and elevated expression on lymphocytes of ITGB7, a RA-modulated gut-homing molecule. ALDH^+^ activity in monocytes positively correlated with coronary plaque burden measured as TPV, LAPV and CAC, with preserved significance after adjustment for FRS and/or HIV-1 parameters. Additionally, ALDH activity in classical monocytes of PWH^+^ART only was linked to pFAI, a measure of pericoronary adipose tissue inflammation linked to cardiovascular risk^75,76^. In contrast, plasma RA levels positively correlated with pFAI in both Pw/oH and PWH^+^ART. These findings suggest that ALDH activity and RA may capture distinct aspects of CVD biology in PWH^+^ART *versus* Pw/oH. This interpretation is consistent with evidence implicating RA signaling in cardiovascular inflammation, endothelial function, and tissue remodeling^57^. Together, these findings point to ALDH activity and retinoid metabolism as contributors to CVD risk in ART-treated PWH.

Age-related alterations in hematopoiesis may contribute to the expansion of ALDH^+^ monocytes in PWH^+^ART. Studies in the REPRIEVE clinical trial demonstrated that clonal hematopoiesis of indeterminate potential (CHIP) occurs in PWH and is associated with cardiovascular risk factors^92^. CHIP-associated mutations can promote inflammatory myeloid phenotypes and atherosclerosis^93,94^. Although CHIP was not assessed in our cohort, clonal hematopoiesis may contribute to the emergence of metabolically unique ALDH^+^ monocytes. Beyond retinoid synthesis, ALDH family members regulate aldehyde detoxification, redox homeostasis, lipid metabolism^53,87^. Thus, ALDH^+^ monocytes may represent a metabolically distinct population, with an ontogeny and contribution to HIV-1 pathogenesis and CVD risk during ART to be elucidated.

Of particular relevance, in our study detectable plasma levels of HIV-1 sgp120 coincided with increased RA concentrations. This is consistent with the presence of RARE in the HIV-1 promoter^60^, the modulation of HIV-1 transcription by RA^59^, thus suggesting that residual viral protein production may be associated with selected aspects of systemic retinoid metabolism. Given previous observations linking sgp120 and markers of HIV-1 persistence to coronary atherosclerosis within the CHACS cohort^46,47^, the relationship between residual viral activity, retinoid metabolism, and CVD warrants further investigation.

Finally, in our CHACS sub-study cohort, statin therapy had minimal impact on immune cell-associated ALDH activity and most retinoid-related biomarkers in the present study. We observed no significant differences in ALDH activity, ITGB7 expression, and plasma RA and chemerin levels in PWH^+^ART receiving or not statins, while plasma RBP4 levels showed a minor increase in statin-treated PWH^+^ART participants. This observation is noteworthy given that the REPRIEVE trial demonstrated a significant reduction in major cardiovascular events with statin therapy in PWH^+^ART^95^. Beyond their lipid-lowering properties, statins exert anti-inflammatory and immunomodulatory effects, including regulation of CCL2/CCR2 axis involved in monocyte recruitment and vascular inflammation^96^. Although interactions between ALDH enzymes and cholesterol metabolism are documented^85,87^, our findings indicate that ALDH-associated pathways may reflect biological processes not adequately targeted by statin therapy, thus explaining their contribution to persistent CVD risk during ART-treated HIV-1 infection^34^.

### Limitations

This study has several limitations. The design of this study does not establish a causal link between increased ALDH activity and CVD risk. Also, it remains unclear whether increased ALDH activity occurs before ART or develops during prolonged treatment as a consequence of residual inflammation and viral activity. Indeed, ART regimens including lamivudine (3TC), may directly induce ALDH activity^97^. Longitudinal studies before/after ART initiation will be required to distinguish between these possibilities. The ALDEFLUOR assay measures ALDH activity and does not distinguish among ALDH isoforms, which exhibit different metabolic functions^53^. Although microbial products can induce ALDH expression and activity in myeloid cells^52^, microbial translocation was not evaluated in this study. The presence/recruitment of ALDH^+^ monocytes into atherosclerotic lesions remains unknown. Contemporary PBMC/plasma samples were not available for all study participants, thus limiting statistical power in correlative studies. The small number of PWH^+^ART receiving statins (n=12) analyzed raises the necessity for caution in result interpretation. Finally, other determinants of retinoid metabolism and CVD risk, including dietary vitamin A intake, microbiome composition, hepatic metabolism, and environmental factors, were not assessed but may contribute to variability in ALDH activity and circulating retinoid levels.

## Conclusion

In conclusion, this study identifies components of the ALDH/RA pathway as novel markers of metabolic/immune dysfunction associated with subclinical coronary atherosclerosis in PWH^+^ART independently of traditional CVD risk factors, HIV-1 parameters, and statin treatment. These findings expand current understanding of the immunological mechanisms associated with CVD pathogenesis during ART-treated HIV-1 infection. Future studies aimed at identifying the predominant ALDH isoforms, characterizing the ontogeny of ALDH^+^ monocytes, and elucidating their contribution to vascular inflammation/atherosclerosis may identify new opportunities for targeted interventions to reduce the CVD risk in PWH on ART.

## Acknowledgments

The authors thank Philippe St Onge, and Dr. Gael Dulude (Flow Cytometry Core Facility, CHUM-Research Center, Montréal, QC, Canada) for expert technical support with polychromatic flow cytometry sorting; Anita Ray for expert technical support with CellEngine algorithms analysis; Olfa Debbeche and Dr. Maria de la Cruz Dominguez Punaro(Biosafety Level 3 Core Facility CHUM-Research Center, Montréal, QC, Canada); Mario Legault (FRQ-S/AIDS and Infectious Diseases Network; Montréal, QC, Canada) for help with ethical approvals and informed consents; Dr. Annie Chamberland, Stéphanie Matte, and Mohamed Sylla for blood processing and managing the cell biobank. Finally, the authors acknowledge the key contribution of all study participants for their crucial contribution to the study.

## Authorship contribution

J.D. performed research, analyzed data, prepared figures, and wrote the paper. M.E-F. designed research, contributed to CHACS biobank building, and wrote the paper. K.B. performed research on pFAI and analyzed data. A.F-M. performed multivariate statistical analysis. M.B. performed research on plasma soluble HIV-1 gp120 and analyzed data. E.M.G., J.M., S.K., and T.R.W.S. helped with PBMCs processing for flow cytometry analysis and generated preliminary results. M.M-P. contributed to CHACS biobank building and management. S.I. and J-P.R. provided access to human samples and clinical information and generated preliminary results. N.C. and A.F. designed research and analyzed data. C.C-L. performed research on coronary atherosclerosis CCTA visualisation/measurement (TPV, LAPV, CAC) and analyzed results. C.T. and M.D., designed research, prepared ethical protocols and written informed consents for the CHACS, provided access to clinical information and build the CHACS biobank. P.A. designed research, analyzed data, prepared figures, and wrote the paper. All authors revised and approved the manuscript.

---

**Supplemental Figure 1 (Related to Figure 2).**
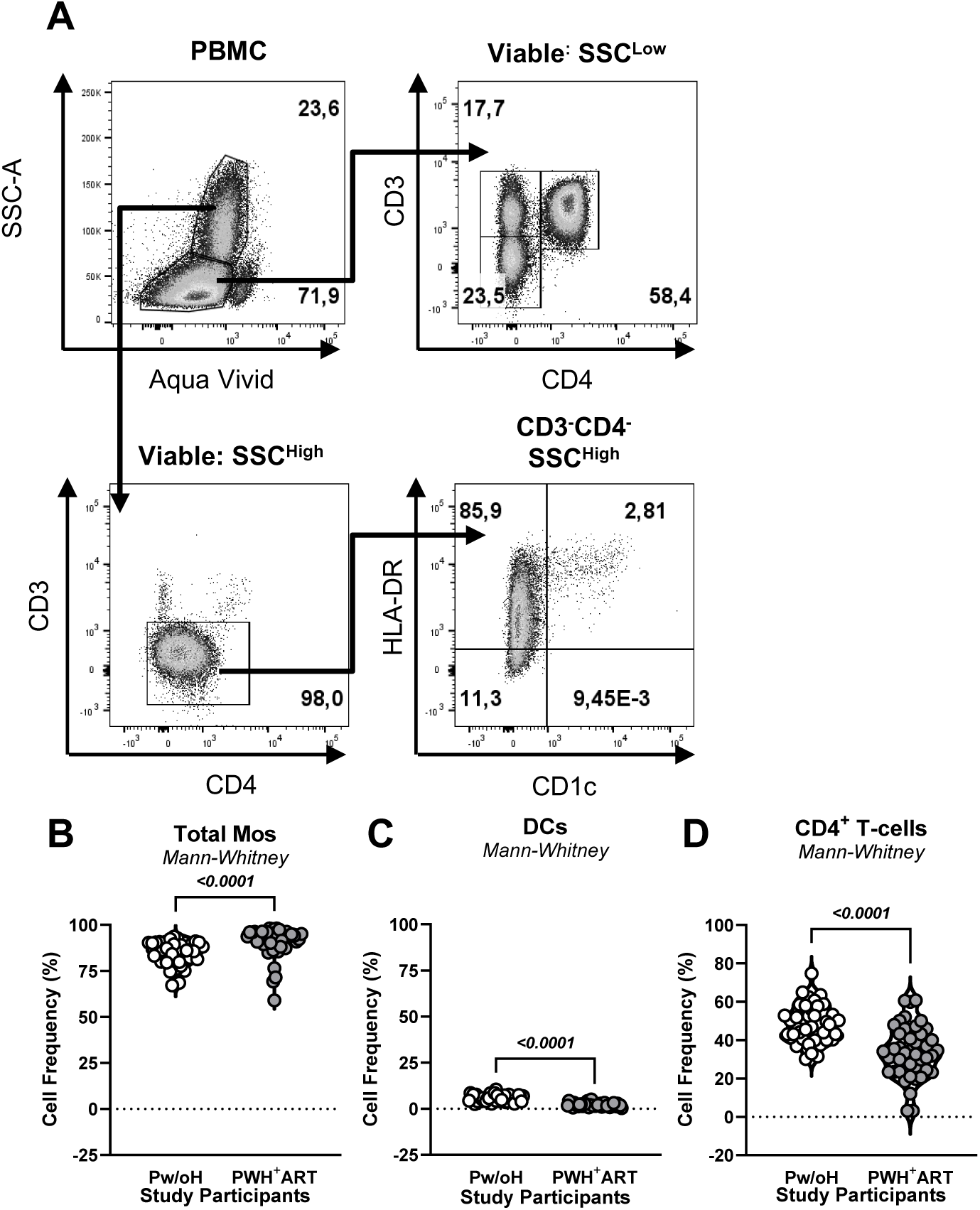
Gating strategy and frequencies of PBMC populations in CHACS participants. **(A)** Representative gating strategy used to identify major PBMC subsets by flow cytometry, including CD3^+^CD4^+^ T-cells, total monocytes, and DC, followed by the identification of classical, intermediate, and non-classical monocyte subsets based on the expression of CD3, CD4, CD1c, HLA-DR, CD14, and CD16. **(B)** Frequencies of total monocytes, **(C)** DC, and **(D)** CD3^+^CD4^+^ T-cells comparing Pw/oH and PWH^+^ART. Statistical analyses were performed using Mann–Whitney tests. P-values are indicated on the graphs.

**Supplemental Figure 2 (Related to Figure 2).**
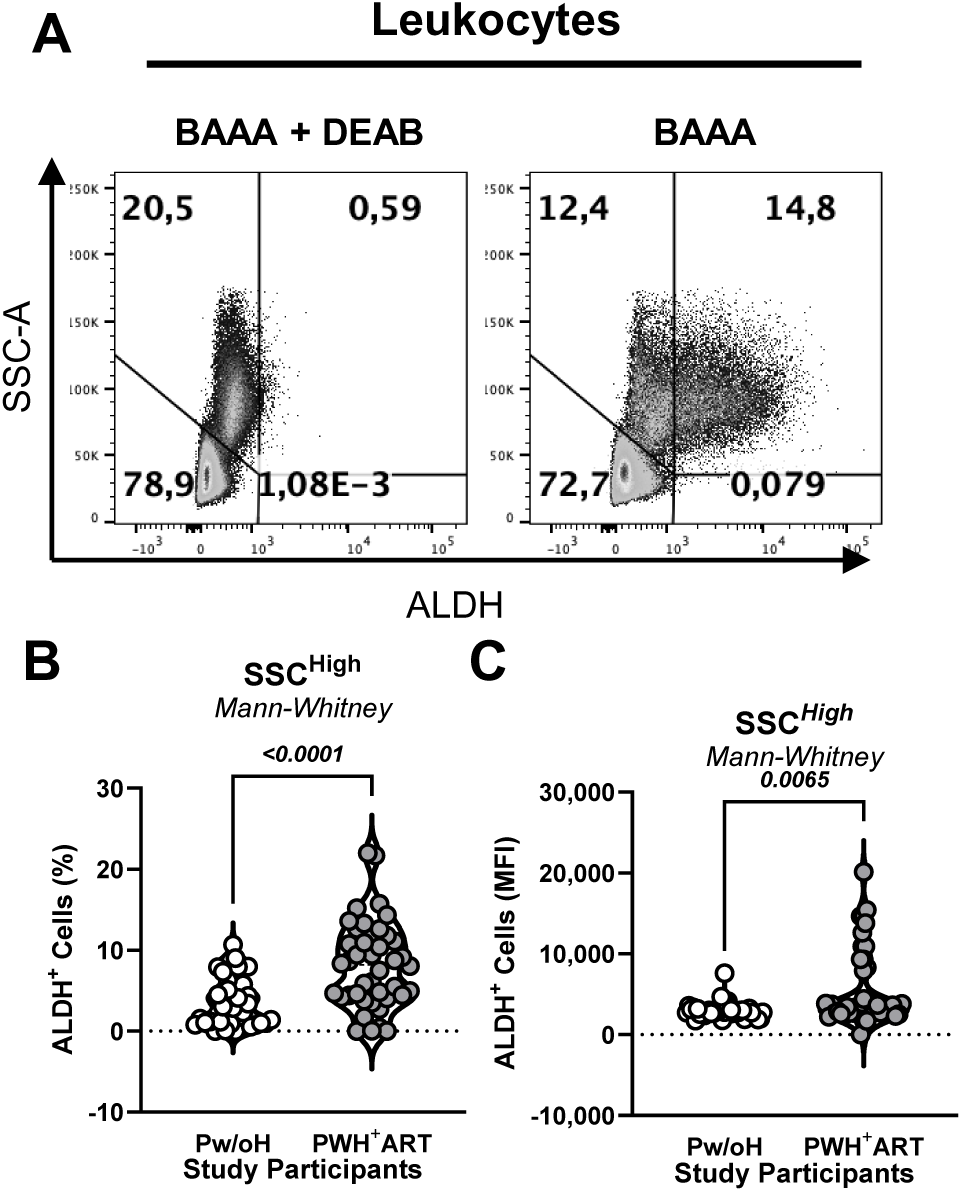
ALDH activity within SSC^High^ leukocyte populations in CHACS participants. **(A)** Representative flow cytometry plots illustrating the gating strategy used to identify ALDH^+^ leukocytes within the SSC^High^ compartment following BAAA staining relative to DEAB-treated controls. **(B)** Frequency of ALDH^+^ SSC^High^ leukocytes (left panel) and MFI of ALDH activity within SSC^High^ leukocytes (right panel) comparing Pw/oH and PWH^+^ART. Statistical analyses were performed using Mann–Whitney tests. P-values are indicated on the graphs.

**Supplemental Figure 3 (Related to Figure 3).**
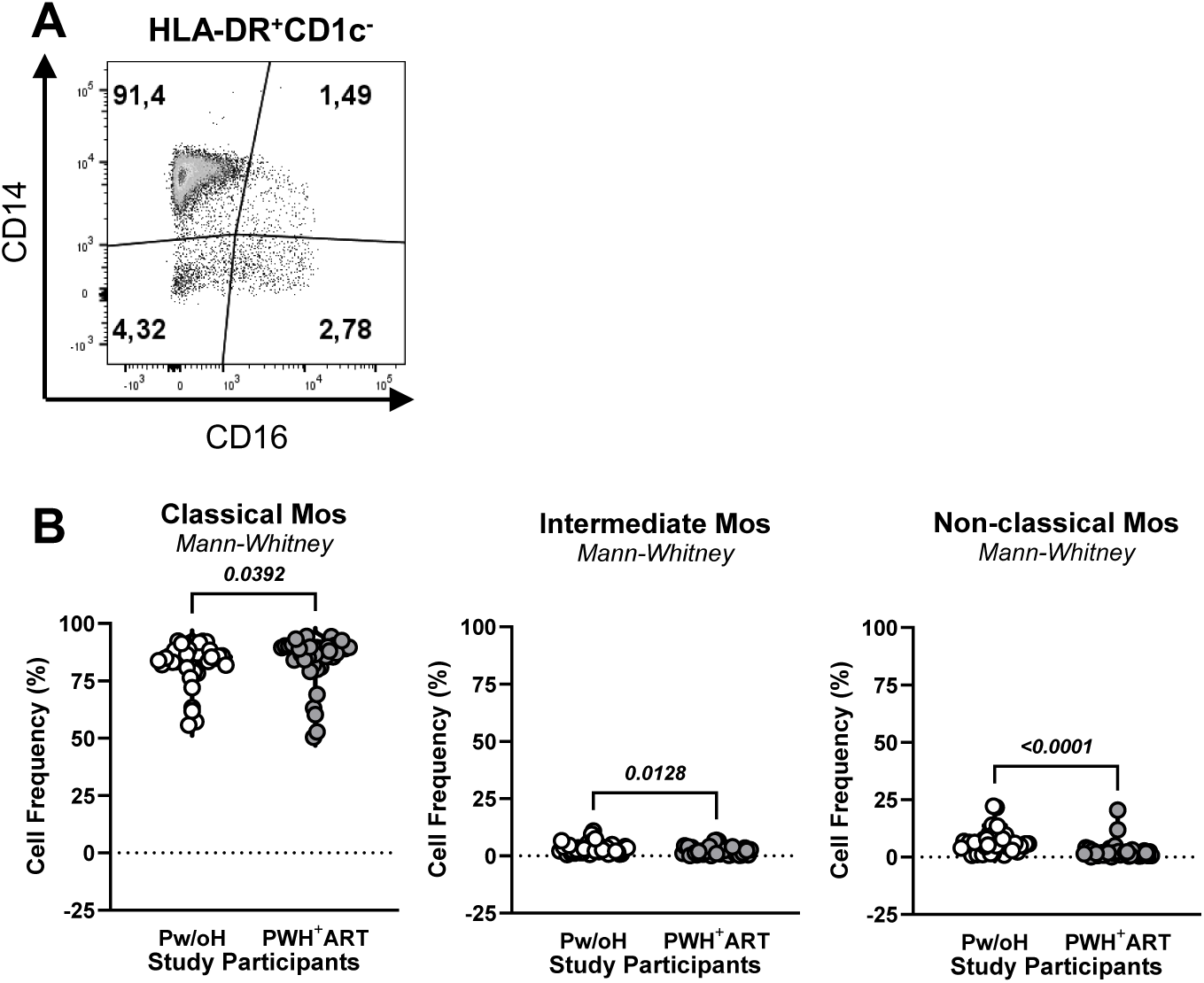
Gating strategy and frequencies of monocyte subsets in CHACS participants. **(A)** Representative gating strategy used to identify classical, intermediate, and non-classical monocyte subsets based on the differential expression of CD14 and CD16 within SSC^High^CD3^−^CD4^−^CD1c^−^HLA-DR^+^ total monocytes. **(B)** Frequencies of classical (left panel), intermediate (middle panel), and non-classical (right panel) monocyte subsets comparing Pw/oH and PWH^+^ART. Statistical analyses were performed using Mann–Whitney tests. P-values are indicated on the graphs.

**Supplemental Figure 4 (Related to Figure 6):**
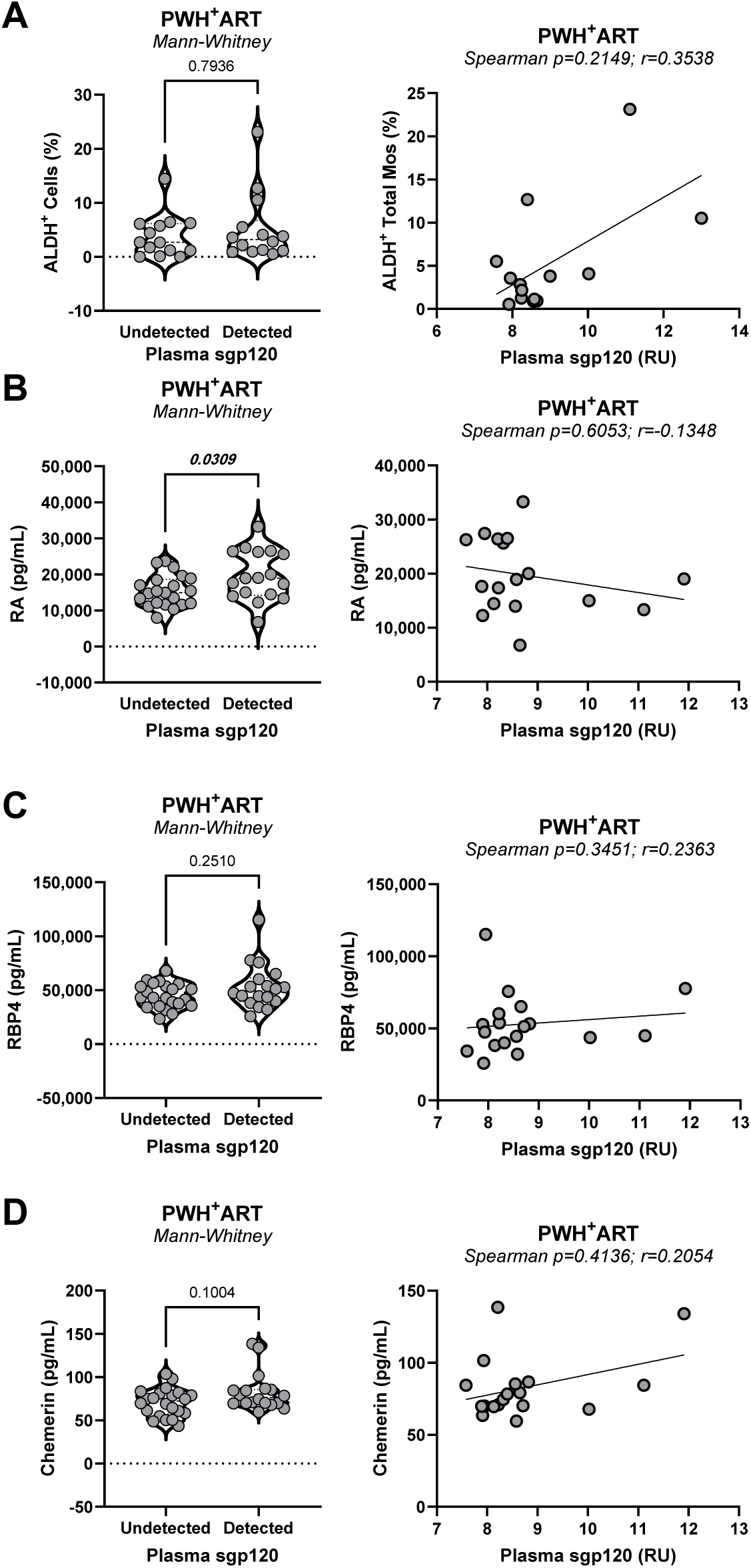
Relationship between sgp120, ALDH activity, and retinoid-associated biomarkers in PWH^+^ART CHACS Participants. **(A)** Frequency of ALDH^+^ total monocytes in PWH^+^ART with detectable or undetectable plasma sgp120 (left panel) and correlation between ALDH^+^ total monocytes and plasma sgp120 levels (right panel). **(B)** Plasma RA concentrations in study participants with detectable or undetectable plasma sgp120 (left panel) and correlation between RA concentrations and plasma sgp120 levels (right panel). **(C)** Plasma RBP4 concentrations in study participants with detectable or undetectable plasma sgp120 (left panel) and correlation between RBP4 concentrations and plasma sgp120 levels (right panel). **(D)** Plasma chemerin concentrations in study participants with detectable or undetectable plasma sgp120 (left panel) and correlation between chemerin concentrations and plasma sgp120 levels (right panel). Two-group comparisons were performed using Mann-Whitney tests, while correlations were assessed using Spearman rank correlation analyses. Statistical p-values and correlation coefficients (r) are indicated on the graphs.

**Supplemental Figure 5 (Related to Figures 2, 3, 4, and 6).**
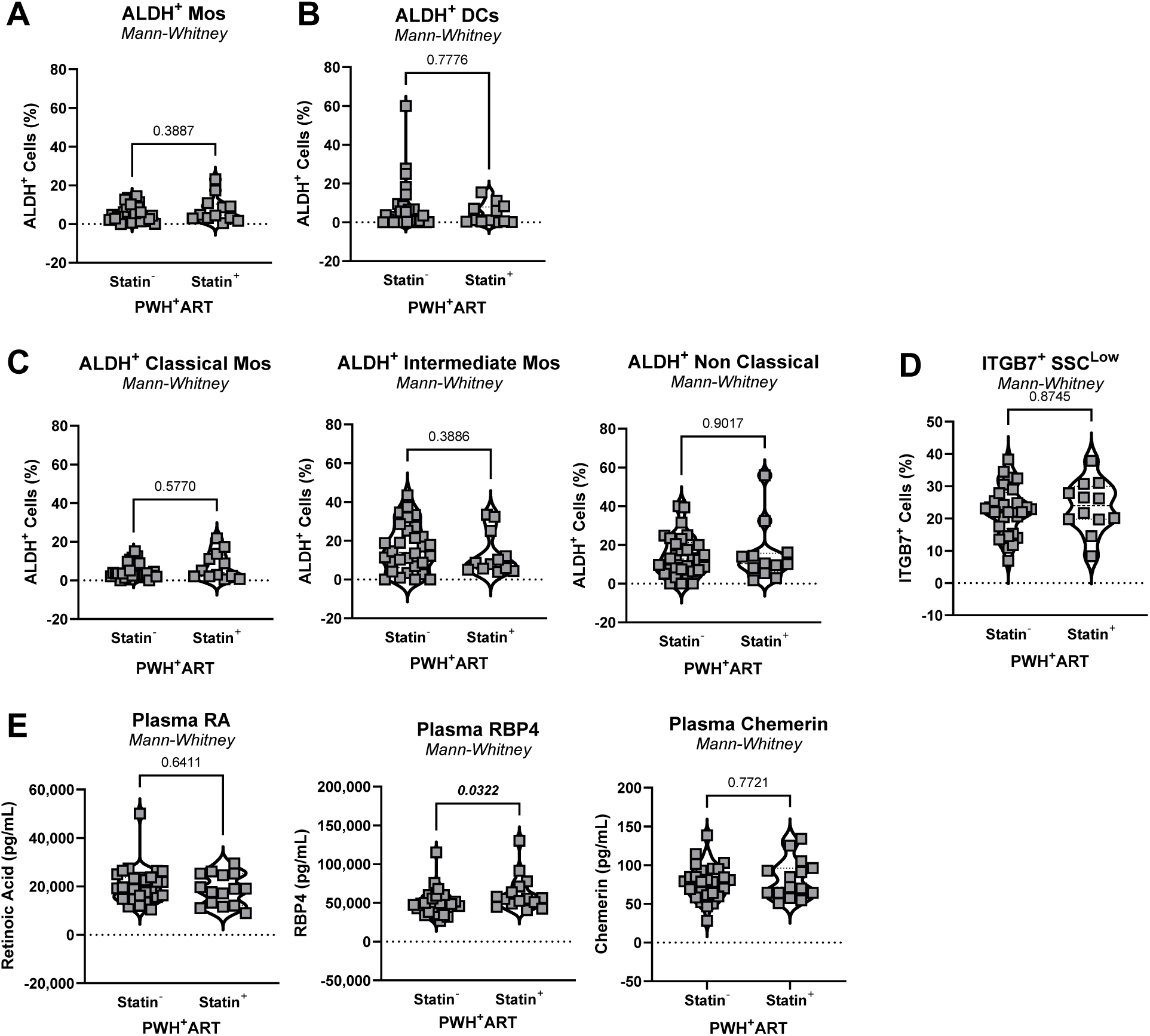
Impact of statin use on ALDH activity, ITGB7 expression, and circulating retinoid-associated biomarkers in PWH+ART CHACS participants. **(A)** Frequencies of ALDH^+^ total monocytes **(B)** DC comparing PWH^+^ART not receiving statin therapy (Statin^−^) and those receiving statin therapy (Statin^+^). **(C)** Frequencies of ALDH^+^ classical (left panel), intermediate (middle panel), and non-classical (right panel) monocyte subsets comparing Statin^−^ and Statin^+^ PWH^+^ART. **(D)** Frequencies of ITGB7^+^ SSC^Low^ leukocyte populations comparing Statin^−^ and Statin^+^ PWH^+^ART. **(E)** Plasma concentrations of RA (left panel), RBP4 (middle panel), and chemerin (right panel) comparing Statin^−^ and Statin^+^ PWH^+^ART. Statistical analyses were performed using Mann–Whitney tests. P-values are indicated on the graphs.

## REFRENCES

1. Bekker, L.G., Beyrer, C., Mgodi, N., Lewin, S.R., Delany-Moretlwe, S., Taiwo, B., Masters, M.C., and Lazarus, J.V. (2023). HIV infection. Nat Rev Dis Primers 9, 42. 10.1038/s41572-023-00452-3.

2. Shin, Y.H., Park, C.M., and Yoon, C.H. (2021). An Overview of Human Immunodeficiency Virus-1 Antiretroviral Drugs: General Principles and Current Status. Infect Chemother 53, 29–45. 10.3947/ic.2020.0100.

3. Deeks, S.G., Archin, N., Cannon, P., Collins, S., Jones, R.B., de Jong, M., Lambotte, O., Lamplough, R., Ndung’u, T., Sugarman, J., et al. (2021). Research priorities for an HIV cure: International AIDS Society Global Scientific Strategy 2021. Nat Med 27, 2085–2098. 10.1038/s41591-021-01590-5.

4. Landovitz, R.J., Scott, H., and Deeks, S.G. (2023). Prevention, treatment and cure of HIV infection. Nat Rev Microbiol 21, 657–670. 10.1038/s41579-023-00914-1.

5. Teer, E., and Essop, M.F. (2018). HIV and Cardiovascular Disease: Role of Immunometabolic Perturbations. Physiology (Bethesda) 33, 74–82. 10.1152/physiol.00028.2017.

6. Siliciano, J.D., and Siliciano, R.F. (2022). In Vivo Dynamics of the Latent Reservoir for HIV-1: New Insights and Implications for Cure. Annu Rev Pathol 17, 271–294. 10.1146/annurev-pathol-050520-112001.

7. Dagand, T., Gigan, J.P., Fromentin, R., and Chomont, N. (2026). The place to be: HIV persistence within tissue reservoirs. Curr Opin HIV AIDS 21, 296–301. 10.1097/COH.0000000000001037.

8. Shah, A.S.V., Stelzle, D., Lee, K.K., Beck, E.J., Alam, S., Clifford, S., Longenecker, C.T., Strachan, F., Bagchi, S., Whiteley, W., et al. (2018). Global Burden of Atherosclerotic Cardiovascular Disease in People Living With HIV: Systematic Review and Meta-Analysis. Circulation 138, 1100–1112. 10.1161/CIRCULATIONAHA.117.033369.

9. Marcus, J.L., Leyden, W.A., Alexeeff, S.E., Anderson, A.N., Hechter, R.C., Hu, H., Lam, J.O., Towner, W.J., Yuan, Q., Horberg, M.A., and Silverberg, M.J. (2020). Comparison of Overall and Comorbidity-Free Life Expectancy Between Insured Adults With and Without HIV Infection, 2000-2016. JAMA Netw Open 3, e207954. 10.1001/jamanetworkopen.2020.7954.

10. Afolabi, J.M., and Kirabo, A. (2024). HIV and Cardiovascular Disease. Circ Res 134, 1512–1514. 10.1161/CIRCRESAHA.124.324805.

11. Feinstein, M.J. (2021). HIV and Cardiovascular Disease: From Insights to Interventions. Top Antivir Med 29, 407–411.

12. Hudson, J.A., Majonga, E.D., Ferrand, R.A., Perel, P., Alam, S.R., and Shah, A.S.V. (2022). Association of HIV Infection With Cardiovascular Pathology Based on Advanced Cardiovascular Imaging: A Systematic Review. JAMA 328, 951–962. 10.1001/jama.2022.15078.

13. Feinstein, M.J. (2022). HIV, Subclinical Cardiovascular Disease, and Clinical Progression: Insights From Immunologic Heterogeneity. JAMA 328, 931–932. 10.1001/jama.2022.15226.

14. Wagner, J.A., Sandel, D.A., Deitchman, A.N., Rutishauser, R.L., Deeks, S.G., and Peluso, M.J. (2026). Advances in Immune-Based Approaches for the Cure of HIV Infection. Drugs 86, 851–870. 10.1007/s40265-026-02311-3.

15. Armani-Tourret, M., Bone, B., Tan, T.S., Sun, W., Bellefroid, M., Struyve, T., Louella, M., Yu, X.G., and Lichterfeld, M. (2024). Immune targeting of HIV-1 reservoir cells: a path to elimination strategies and cure. Nat Rev Microbiol 22, 328–344. 10.1038/s41579-024-01010-8.

16. Ijaz, A., Rasheed, A., and Orecchioni, M. (2026). Macrophages in human atherosclerotic plaques in the era of single-cell and spatial transcriptomics. Immunohorizons 10. 10.1093/immhor/vlaf089.

17. Chen, R., Zhang, H., Tang, B., Luo, Y., Yang, Y., Zhong, X., Chen, S., Xu, X., Huang, S., and Liu, C. (2024). Macrophages in cardiovascular diseases: molecular mechanisms and therapeutic targets. Signal Transduct Target Ther 9, 130. 10.1038/s41392-024-01840-1.

18. Gupta, N., Liu, Y., and Cai, B. (2026). Macrophage efferocytosis in cardiovascular disease: mechanisms and therapeutic implications. Immunohorizons 10. 10.1093/immhor/vlag002.

19. Ajasin, D., Arredondo-Anez, S., Gutierrez, J.A., Ward, M.C., Maass, K., and Eugenin, E. (2026). HIV heart inflammation is mediated by HIV infected myeloid cells, HIV-tat secretion, and aberrant function of Connexin43-containing channels. Sci Rep 16. 10.1038/s41598-026-43625-2.

20. Winkels, H., Ehinger, E., Vassallo, M., Buscher, K., Dinh, H.Q., Kobiyama, K., Hamers, A.A.J., Cochain, C., Vafadarnejad, E., Saliba, A.E., et al. (2018). Atlas of the Immune Cell Repertoire in Mouse Atherosclerosis Defined by Single-Cell RNA-Sequencing and Mass Cytometry. Circ Res 122, 1675–1688. 10.1161/CIRCRESAHA.117.312513.

21. Fernandez, D.M., Rahman, A.H., Fernandez, N.F., Chudnovskiy, A., Amir, E.D., Amadori, L., Khan, N.S., Wong, C.K., Shamailova, R., Hill, C.A., et al. (2019). Single-cell immune landscape of human atherosclerotic plaques. Nat Med 25, 1576–1588. 10.1038/s41591-019-0590-4.

22. Roy, P., Orecchioni, M., and Ley, K. (2022). How the immune system shapes atherosclerosis: roles of innate and adaptive immunity. Nat Rev Immunol 22, 251–265. 10.1038/s41577-021-00584-1.

23. Tabas, I., and Bornfeldt, K.E. (2016). Macrophage Phenotype and Function in Different Stages of Atherosclerosis. Circ Res 118, 653–667. 10.1161/CIRCRESAHA.115.306256.

24. Cochain, C., Vafadarnejad, E., Arampatzi, P., Pelisek, J., Winkels, H., Ley, K., Wolf, D., Saliba, A.E., and Zernecke, A. (2018). Single-Cell RNA-Seq Reveals the Transcriptional Landscape and Heterogeneity of Aortic Macrophages in Murine Atherosclerosis. Circ Res 122, 1661–1674. 10.1161/CIRCRESAHA.117.312509.

25. Cybulsky, M.I., Cheong, C., and Robbins, C.S. (2016). Macrophages and Dendritic Cells: Partners in Atherogenesis. Circ Res 118, 637–652. 10.1161/CIRCRESAHA.115.306542.

26. Meng, X., Yang, J., Dong, M., Zhang, K., Tu, E., Gao, Q., Chen, W., Zhang, C., and Zhang, Y. (2016). Regulatory T cells in cardiovascular diseases. Nat Rev Cardiol 13, 167–179. 10.1038/nrcardio.2015.169.

27. Yun, T.J., Lee, J.S., Machmach, K., Shim, D., Choi, J., Wi, Y.J., Jang, H.S., Jung, I.H., Kim, K., Yoon, W.K., et al. (2016). Indoleamine 2,3-Dioxygenase-Expressing Aortic Plasmacytoid Dendritic Cells Protect against Atherosclerosis by Induction of Regulatory T Cells. Cell Metab 23, 852–866. 10.1016/j.cmet.2016.04.010.

28. D’Agostino, R.B., Sr., Vasan, R.S., Pencina, M.J., Wolf, P.A., Cobain, M., Massaro, J.M., and Kannel, W.B. (2008). General cardiovascular risk profile for use in primary care: the Framingham Heart Study. Circulation 117, 743–753. 10.1161/CIRCULATIONAHA.107.699579.

29. Triant, V.A., Lyass, A., Hurley, L.B., Borowsky, L.H., Ehrbar, R.Q., He, W., Cheng, D., Lo, J., Klein, D.B., Meigs, J.B., et al. (2024). Cardiovascular Risk Estimation Is Suboptimal in People With HIV. J Am Heart Assoc 13, e029228. 10.1161/JAHA.123.029228.

30. Zou, R.S., Ruan, Y., Truong, B., Bhattacharya, R., Lu, M.T., Karady, J., Bernardo, R., Finneran, P., Hornsby, W., Fitch, K.V., et al. (2024). Polygenic Scores and Preclinical Cardiovascular Disease in Individuals With HIV: Insights From the REPRIEVE Trial. J Am Heart Assoc 13, e033413. 10.1161/JAHA.123.033413.

31. Spray, L., Richardson, G., Haendeler, J., Altschmied, J., Rumampouw, V., Wallis, S.B., Georgiopoulos, G., White, S., Unsworth, A., Stellos, K., et al. (2025). Cardiovascular inflammaging: Mechanisms, consequences, and therapeutic perspectives. Cell Rep Med 6, 102264. 10.1016/j.xcrm.2025.102264.

32. Ntsekhe, M., and Baker, J.V. (2023). Cardiovascular Disease Among Persons Living With HIV: New Insights Into Pathogenesis and Clinical Manifestations in a Global Context. Circulation 147, 83–100. 10.1161/CIRCULATIONAHA.122.057443.

33. Ghandakly, E., Moudgil, R., and Holman, K. (2025). Cardiovascular disease in people living with HIV: Risk assessment and management. Cleve Clin J Med 92, 159–167. 10.3949/ccjm.92a.24055.

34. Obare, L.M., Stephens, V.R., and Wanjalla, C.N. (2025). Understanding residual risk of cardiovascular disease in people with HIV. Curr Opin HIV AIDS 20, 319–330. 10.1097/COH.0000000000000953.

35. Feinstein, M.J., Hsue, P.Y., Benjamin, L.A., Bloomfield, G.S., Currier, J.S., Freiberg, M.S., Grinspoon, S.K., Levin, J., Longenecker, C.T., and Post, W.S. (2019). Characteristics, Prevention, and Management of Cardiovascular Disease in People Living With HIV: A Scientific Statement From the American Heart Association. Circulation 140, e98–e124. 10.1161/CIR.0000000000000695.

36. Yoshino, Y., Kimura, Y., and Ito, F. (2026). Chronic inflammation in virus-suppressed people living with human immunodeficiency virus infection: A microbiology-oriented perspective on gut barrier failure, microbial translocation, and immune activation. J Infect Chemother 32, 102972. 10.1016/j.jiac.2026.102972.

37. Deeks, S.G., Lewin, S.R., Ross, A.L., Ananworanich, J., Benkirane, M., Cannon, P., Chomont, N., Douek, D., Lifson, J.D., Lo, Y.R., et al. (2016). International AIDS Society global scientific strategy: towards an HIV cure 2016. Nat Med 22, 839–850. 10.1038/nm.4108.

38. Siliciano, J.D., and Siliciano, R.F. (2014). Recent developments in the search for a cure for HIV-1 infection: targeting the latent reservoir for HIV-1. J Allergy Clin Immunol 134, 12–19. 10.1016/j.jaci.2014.05.026.

39. Chun, T.W., and Fauci, A.S. (2012). HIV reservoirs: pathogenesis and obstacles to viral eradication and cure. AIDS 26, 1261–1268. 10.1097/QAD.0b013e328353f3f1.

40. Finzi, D., Hermankova, M., Pierson, T., Carruth, L.M., Buck, C., Chaisson, R.E., Quinn, T.C., Chadwick, K., Margolick, J., Brookmeyer, R., et al. (1997). Identification of a reservoir for HIV-1 in patients on highly active antiretroviral therapy. Science 278, 1295–1300. 10.1126/science.278.5341.1295.

41. Hatano, H., Delwart, E.L., Norris, P.J., Lee, T.H., Neilands, T.B., Kelley, C.F., Hunt, P.W., Hoh, R., Linnen, J.M., Martin, J.N., et al. (2010). Evidence of persistent low-level viremia in long-term HAART-suppressed, HIV-infected individuals. AIDS 24, 2535–2539. 10.1097/QAD.0b013e32833dba03.

42. McLaughlin, M.M., Ma, Y., Scherzer, R., Rahalkar, S., Martin, J.N., Mills, C., Milush, J., Deeks, S.G., and Hsue, P.Y. (2020). Association of Viral Persistence and Atherosclerosis in Adults With Treated HIV Infection. JAMA Netw Open 3, e2018099. 10.1001/jamanetworkopen.2020.18099.

43. Hsue, P.Y., and Waters, D.D. (2019). HIV infection and coronary heart disease: mechanisms and management. Nat Rev Cardiol 16, 745–759. 10.1038/s41569-019-0219-9.

44. Wang, Y., Brichacek, B., Dubrovsky, L., Pushkarsky, T., Korolowicz, K., Rodriguez, O., Lee, Y., Catalfamo, M., Albanese, C., Popratiloff, A., et al. (2025). Increased atherosclerosis in HIV-infected humanized mice is caused by a single viral protein, Nef. J Infect Dis. 10.1093/infdis/jiaf192.

45. Hudson, J.A., Ferrand, R.A., Gitau, S.N., Mureithi, M.W., Maffia, P., Alam, S.R., and Shah, A.S.V. (2024). HIV-Associated Cardiovascular Disease Pathogenesis: An Emerging Understanding Through Imaging and Immunology. Circ Res 134, 1546–1565. 10.1161/CIRCRESAHA.124.323890.

46. Turcotte, I., El-Far, M., Sadouni, M., Chartrand-Lefebvre, C., Filali-Mouhim, A., Fromentin, R., Chamberland, A., Jenabian, M.A., Baril, J.G., Trottier, B., et al. (2023). Association Between the Development of Subclinical Cardiovascular Disease and Human Immunodeficiency Virus (HIV) Reservoir Markers in People With HIV on Suppressive Antiretroviral Therapy. Clin Infect Dis 76, 1318–1321. 10.1093/cid/ciac874.

47. Benlarbi, M., Richard, J., Bourassa, C., Tolbert, W.D., Chartrand-Lefebvre, C., Gendron-Lepage, G., Sylla, M., El-Far, M., Messier-Peet, M., Guertin, C., et al. (2024). Plasma Human Immunodeficiency Virus 1 Soluble Glycoprotein 120 Association With Correlates of Immune Dysfunction and Inflammation in Antiretroviral Therapy-Treated Individuals With Undetectable Viremia. J Infect Dis 229, 763–774. 10.1093/infdis/jiad503.

48. Ramani, H., Gosselin, A., Bunet, R., Jenabian, M.A., Sylla, M., Pagliuzza, A., Chartrand-Lefebvre, C., Routy, J.P., Goulet, J.P., Thomas, R., et al. (2024). IL-32 Drives the Differentiation of Cardiotropic CD4+ T Cells Carrying HIV DNA in People With HIV. J Infect Dis 229, 1277–1289. 10.1093/infdis/jiad576.

49. Brenchley, J.M., and Douek, D.C. (2012). Microbial translocation across the GI tract. Annu Rev Immunol 30, 149–173. 10.1146/annurev-immunol-020711-075001.

50. Nganou-Makamdop, K., and Douek, D.C. (2024). The Gut and the Translocated Microbiomes in HIV Infection: Current Concepts and Future Avenues. Pathog Immun 9, 168–194. 10.20411/pai.v9i1.693.

51. Planas, D., Routy, J.P., and Ancuta, P. (2019). New Th17-specific therapeutic strategies for HIV remission. Curr Opin HIV AIDS 14, 85–92. 10.1097/COH.0000000000000522.

52. Cattin, A., Wacleche, V.S., Fonseca Do Rosario, N., Marchand, L.R., Dias, J., Gosselin, A., Cohen, E.A., Estaquier, J., Chomont, N., Routy, J.P., and Ancuta, P. (2021). RALDH Activity Induced by Bacterial/Fungal Pathogens in CD16(+) Monocyte-Derived Dendritic Cells Boosts HIV Infection and Outgrowth in CD4(+) T Cells. J Immunol 206, 2638–2651. 10.4049/jimmunol.2001436.

53. Zhao, W., Xia, Y., Gao, Z., Chen, J., and Zhang, E. (2025). Multiple roles of ALDH1 in health and disease. Front Physiol 16, 1627164. 10.3389/fphys.2025.1627164.

54. Iwata, M., Hirakiyama, A., Eshima, Y., Kagechika, H., Kato, C., and Song, S.Y. (2004). Retinoic acid imprints gut-homing specificity on T cells. Immunity 21, 527–538. 10.1016/j.immuni.2004.08.011.

55. Bakdash, G., Vogelpoel, L.T., van Capel, T.M., Kapsenberg, M.L., and de Jong, E.C. (2015). Retinoic acid primes human dendritic cells to induce gut-homing, IL-10-producing regulatory T cells. Mucosal Immunol 8, 265–278. 10.1038/mi.2014.64.

56. Wacleche, V.S., Cattin, A., Goulet, J.P., Gauchat, D., Gosselin, A., Cleret-Buhot, A., Zhang, Y., Tremblay, C.L., Routy, J.P., and Ancuta, P. (2018). CD16(+) monocytes give rise to CD103(+)RALDH2(+)TCF4(+) dendritic cells with unique transcriptional and immunological features. Blood Adv 2, 2862–2878. 10.1182/bloodadvances.2018020123.

57. Parker, L.E., Papanicolaou, K.N., Zalesak-Kravec, S., Weinberger, E.M., Kane, M.A., and Foster, D.B. (2025). Retinoic acid signaling and metabolism in heart failure. Am J Physiol Heart Circ Physiol 328, H792–H813. 10.1152/ajpheart.00871.2024.

58. Planas, D., Zhang, Y., Monteiro, P., Goulet, J.P., Gosselin, A., Grandvaux, N., Hope, T.J., Fassati, A., Routy, J.P., and Ancuta, P. (2017). HIV-1 selectively targets gut-homing CCR6+CD4+ T cells via mTOR-dependent mechanisms. JCI Insight 2. 10.1172/jci.insight.93230.

59. Dias, J., Cattin, A., Bendoumou, M., Dutilleul, A., Lodge, R., Goulet, J.P., Fert, A., Raymond Marchand, L., Wiche Salinas, T.R., Ngassaki Yoka, C.D., et al. (2024). Retinoic acid enhances HIV-1 reverse transcription and transcription in macrophages via mTOR-modulated mechanisms. Cell Rep 43, 114414. 10.1016/j.celrep.2024.114414.

60. Lee, M.O., Hobbs, P.D., Zhang, X.K., Dawson, M.I., and Pfahl, M. (1994). A synthetic retinoid antagonist inhibits the human immunodeficiency virus type 1 promoter. Proc Natl Acad Sci U S A 91, 5632–5636. 10.1073/pnas.91.12.5632.

61. Rhee, E.J., Nallamshetty, S., and Plutzky, J. (2012). Retinoid metabolism and its effects on the vasculature. Biochim Biophys Acta 1821, 230–240. 10.1016/j.bbalip.2011.07.001.

62. Correction to: Association of Serum Retinoic Acid With Risk of Mortality in Patients With Coronary Artery Disease. (2017). Circ Res 121, e84. 10.1161/RES.0000000000000180.

63. Bilbija, D., Elmabsout, A.A., Sagave, J., Haugen, F., Bastani, N., Dahl, C.P., Gullestad, L., Sirsjo, A., Blomhoff, R., and Valen, G. (2014). Expression of retinoic acid target genes in coronary artery disease. Int J Mol Med 33, 677–686. 10.3892/ijmm.2014.1623.

64. Graham, T.E., Yang, Q., Bluher, M., Hammarstedt, A., Ciaraldi, T.P., Henry, R.R., Wason, C.J., Oberbach, A., Jansson, P.A., Smith, U., and Kahn, B.B. (2006). Retinol-binding protein 4 and insulin resistance in lean, obese, and diabetic subjects. N Engl J Med 354, 2552–2563. 10.1056/NEJMoa054862.

65. Sun, Q., Kiernan, U.A., Shi, L., Phillips, D.A., Kahn, B.B., Hu, F.B., Manson, J.E., Albert, C.M., and Rexrode, K.M. (2013). Plasma retinol-binding protein 4 (RBP4) levels and risk of coronary heart disease: a prospective analysis among women in the nurses’ health study. Circulation 127, 1938–1947. 10.1161/CIRCULATIONAHA.113.002073.

66. Pastorio, C., Richard, K., Usmani, S., Kissmann, A.K., Bolotnikov, G., Gosalbez, G., Hayn, M., Koepke, L., Sauertnik, A., Preising, A., et al. (2025). Retinol Binding Protein 4 reactivates latent HIV-1 by triggering canonical NF-kappaB, JAK/STAT5 and JNK signalling. Signal Transduct Target Ther 10, 326. 10.1038/s41392-025-02424-3.

67. Bozaoglu, K., Bolton, K., McMillan, J., Zimmet, P., Jowett, J., Collier, G., Walder, K., and Segal, D. (2007). Chemerin is a novel adipokine associated with obesity and metabolic syndrome. Endocrinology 148, 4687–4694. 10.1210/en.2007-0175.

68. Ernst, M.C., and Sinal, C.J. (2010). Chemerin: at the crossroads of inflammation and obesity. Trends Endocrinol Metab 21, 660–667. 10.1016/j.tem.2010.08.001.

69. Durand, M., Chartrand-Lefebvre, C., Baril, J.G., Trottier, S., Trottier, B., Harris, M., Walmsley, S., Conway, B., Wong, A., Routy, J.P., et al. (2017). The Canadian HIV and aging cohort study - determinants of increased risk of cardio-vascular diseases in HIV-infected individuals: rationale and study protocol. BMC Infect Dis 17, 611. 10.1186/s12879-017-2692-2.

70. Giguere, K., Chartrand-Lefebvre, C., Baril, J.G., Conway, B., El-Far, M., Falutz, J., Harris, M., Jenabian, M.A., Leipsic, J., Loutfy, M., et al. (2023). Baseline characteristics of a prospective cohort study of aging and cardiovascular diseases among people living with HIV. HIV Med 24, 1210–1221. 10.1111/hiv.13550.

71. Boldeanu, I., Sadouni, M., Mansour, S., Baril, J.G., Trottier, B., Soulez, G., A, S.C., Leipsic, J., Tremblay, C., Durand, M., et al. (2021). Prevalence and Characterization of Subclinical Coronary Atherosclerotic Plaque with CT among Individuals with HIV: Results from the Canadian HIV and Aging Cohort Study. Radiology 299, 571–580. 10.1148/radiol.2021203297.

72. Wiche Salinas, T.R., Zhang, Y., Gosselin, A., Rosario, N.F., El-Far, M., Filali-Mouhim, A., Routy, J.P., Chartrand-Lefebvre, C., Landay, A.L., Durand, M., et al. (2024). Alterations in Th17 Cells and Non-Classical Monocytes as a Signature of Subclinical Coronary Artery Atherosclerosis during ART-Treated HIV-1 Infection. Cells 13. 10.3390/cells13020157.

73. Moreira Gabriel, E., Dias, J., Filali-Mouhim, A., Caballero, R.E., Wiche Salinas, T.R., Nayrac, M., Chartrand-Lefebvre, C., Routy, J.P., Durand, M., El-Far, M., et al. (2025). Alterations in Circulating T-Cell Subsets with Gut-Homing/Residency Phenotypes Associated with HIV-1 Status and Subclinical Atherosclerosis. Cells 14. 10.3390/cells14211732.

74. Chen, Z., Boldeanu, I., Nepveu, S., Durand, M., Chin, A.S., Kauffmann, C., Mansour, S., Soulez, G., Tremblay, C., and Chartrand-Lefebvre, C. (2017). In vivo coronary artery plaque assessment with computed tomography angiography: is there an impact of iterative reconstruction on plaque volume and attenuation metrics? Acta Radiol 58, 660–669. 10.1177/0284185116664229.

75. Antonopoulos, A.S., Sanna, F., Sabharwal, N., Thomas, S., Oikonomou, E.K., Herdman, L., Margaritis, M., Shirodaria, C., Kampoli, A.M., Akoumianakis, I., et al. (2017). Detecting human coronary inflammation by imaging perivascular fat. Sci Transl Med 9. 10.1126/scitranslmed.aal2658.

76. Antoniades, C., Antonopoulos, A.S., and Deanfield, J. (2020). Imaging residual inflammatory cardiovascular risk. Eur Heart J 41, 748–758. 10.1093/eurheartj/ehz474.

77. Roederer, M. (2002). Compensation in flow cytometry. Curr Protoc Cytom Chapter 1, Unit 1 14. 10.1002/0471142956.cy0114s22.

78. Kenward, M.G., and Roger, J.H. (2010). The use of baseline covariates in crossover studies. Biostatistics 11, 1–17. 10.1093/biostatistics/kxp046.

79. Motoyama, S., Sarai, M., Harigaya, H., Anno, H., Inoue, K., Hara, T., Naruse, H., Ishii, J., Hishida, H., Wong, N.D., et al. (2009). Computed tomographic angiography characteristics of atherosclerotic plaques subsequently resulting in acute coronary syndrome. J Am Coll Cardiol 54, 49–57. 10.1016/j.jacc.2009.02.068.

80. Budoff, M.J., Shaw, L.J., Liu, S.T., Weinstein, S.R., Mosler, T.P., Tseng, P.H., Flores, F.R., Callister, T.Q., Raggi, P., and Berman, D.S. (2007). Long-term prognosis associated with coronary calcification: observations from a registry of 25,253 patients. J Am Coll Cardiol 49, 1860–1870. 10.1016/j.jacc.2006.10.079.

81. Liu, J., and Wang, Y. (2025). The Potential Role of Retinol-Binding Protein 4 in Heart Failure: A Review. Rev Cardiovasc Med 26, 40127. 10.31083/RCM40127.

82. Steinhoff, J.S., Lass, A., and Schupp, M. (2022). Retinoid Homeostasis and Beyond: How Retinol Binding Protein 4 Contributes to Health and Disease. Nutrients 14. 10.3390/nu14061236.

83. Tan, L., Lu, X., Danser, A.H.J., and Verdonk, K. (2023). The Role of Chemerin in Metabolic and Cardiovascular Disease: A Literature Review of Its Physiology and Pathology from a Nutritional Perspective. Nutrients 15. 10.3390/nu15132878.

84. McMillan, R., and Kirabo, A. (2025). Chemerin as a Mediator of Hypertension and Cardiometabolic Diseases (A Comprehensive Review). Curr Hypertens Rep 28, 4. 10.1007/s11906-025-01354-3.

85. Kato, S., Liberona, M.F., Cerda-Infante, J., Sanchez, M., Henriquez, J., Bizama, C., Bravo, M.L., Gonzalez, P., Gejman, R., Branes, J., et al. (2018). Simvastatin interferes with cancer ‘stem-cell’ plasticity reducing metastasis in ovarian cancer. Endocr Relat Cancer 25, 821–836. 10.1530/ERC-18-0132.

86. Li, L., Zhong, S., Li, R., Liang, N., Zhang, L., Xia, S., Xu, X., Chen, X., Chen, S., Tao, Y., and Yin, H. (2022). Aldehyde dehydrogenase 2 and PARP1 interaction modulates hepatic HDL biogenesis by LXRalpha-mediated ABCA1 expression. JCI Insight 7. 10.1172/jci.insight.155869.

87. Zhong, S., Li, L., Liang, N., Zhang, L., Xu, X., Chen, S., and Yin, H. (2021). Acetaldehyde Dehydrogenase 2 regulates HMG-CoA reductase stability and cholesterol synthesis in the liver. Redox Biol 41, 101919. 10.1016/j.redox.2021.101919.

88. Bui, T.V.A., Hwangbo, H., Lai, Y., Hong, S.B., Choi, Y.J., Park, H.J., and Ban, K. (2023). The Gut-Heart Axis: Updated Review for The Roles of Microbiome in Cardiovascular Health. Korean Circ J 53, 499–518. 10.4070/kcj.2023.0048.

89. Ouyang, J., Yan, J., Zhou, X., Isnard, S., Harypursat, V., Cui, H., Routy, J.P., and Chen, Y. (2023). Relevance of biomarkers indicating gut damage and microbial translocation in people living with HIV. Front Immunol 14, 1173956. 10.3389/fimmu.2023.1173956.

90. Wang, Z., Peters, B.A., Bryant, M., Hanna, D.B., Schwartz, T., Wang, T., Sollecito, C.C., Usyk, M., Grassi, E., Wiek, F., et al. (2023). Gut microbiota, circulating inflammatory markers and metabolites, and carotid artery atherosclerosis in HIV infection. Microbiome 11, 119. 10.1186/s40168-023-01566-2.

91. Peters, B.A., Burk, R.D., Kaplan, R.C., and Qi, Q. (2023). The Gut Microbiome, Microbial Metabolites, and Cardiovascular Disease in People Living with HIV. Curr HIV/AIDS Rep 20, 86–99. 10.1007/s11904-023-00648-y.

92. Bhattacharya, R., Uddin, M.M., Patel, A.P., Niroula, A., Finneran, P., Bernardo, R., Fitch, K.V., Lu, M.T., Bloomfield, G.S., Malvestutto, C., et al. (2024). Risk factors for clonal hematopoiesis of indeterminate potential in people with HIV: a report from the REPRIEVE trial. Blood Adv 8, 959–967. 10.1182/bloodadvances.2023011324.

93. Jaiswal, S., Natarajan, P., Silver, A.J., Gibson, C.J., Bick, A.G., Shvartz, E., McConkey, M., Gupta, N., Gabriel, S., Ardissino, D., et al. (2017). Clonal Hematopoiesis and Risk of Atherosclerotic Cardiovascular Disease. 377, 111–121. 10.1056/NEJMoa1701719.

94. Fuster, J.J., MacLauchlan, S., Zuriaga, M.A., Polackal, M.N., Ostriker, A.C., Chakraborty, R., Wu, C.L., Sano, S., Muralidharan, S., Rius, C., et al. (2017). Clonal hematopoiesis associated with TET2 deficiency accelerates atherosclerosis development in mice. Science 355, 842–847. 10.1126/science.aag1381.

95. Grinspoon, S.K., Fitch, K.V., Zanni, M.V., Fichtenbaum, C.J., Umbleja, T., Aberg, J.A., Overton, E.T., Malvestutto, C.D., Bloomfield, G.S., Currier, J.S., et al. (2023). Pitavastatin to Prevent Cardiovascular Disease in HIV Infection. N Engl J Med 389, 687–699. 10.1056/NEJMoa2304146.

96. Gholamalizadeh, H., Ensan, B., Karav, S., Jamialahmadi, T., and Sahebkar, A. (2024). Regulatory effects of statins on CCL2/CCR2 axis in cardiovascular diseases: new insight into pleiotropic effects of statins. J Inflamm (Lond) 21, 51. 10.1186/s12950-024-00420-y.

97. Han, L., Zhou, J., Hu, Z., Fu, C., Li, X., Liu, J., Zhao, W., Wu, T., Li, C., Kang, J., et al. (2022). Lamivudine remedies alcoholism by activating acetaldehyde dehydrogenase. Biochem Pharmacol 203, 115199. 10.1016/j.bcp.2022.115199.

