## Supplemental Table 1 for "ALDH Activity in Monocytes is Associated with Subclinical Coronary Atherosclerosis in Treated People with HIV-1"

**Supplemental Table 1: Key Ressources**

| | SOURCE | IDENTIFIER |
|---|---|---|
| **Leukapheresis** | | |
| Lymphocyte Separation Medium (LSM) | Wisent | Cat#305-010-CL |
| Fetal Bovine Serum (FBS) | Wisent | Cat#091-150 |
| Trypan Blue | Thermo Fisher | Cat#15250061 |
| Dimethyl Sulfoxide (DMSO) | Sigma | Cat#34869-500mL |
| RPMI 1640 Medium (RPMI) | Thermo Fisher | Cat#11875119 |
| Penicillin/streptomycin | Thermo Fisher | Cat#15140122 |
| **Flow cytometry** | | |
| Mouse anti-human CD3 Pacific Blue (Clone UCHT1) | BD | Cat#558117 |
| Mouse anti-human CD4 Alexa Fluor 700 (Clone RPA-T4) | BD | Cat#557922 |
| Mouse anti-human CD16 Phycoerythrin-Cyanine 7 (Clone 3G8) | BD | Cat#560918 |
| Mouse anti-human CD14 APC (Clone M5E2) | BD | Cat#555399 |
| Anti-human CD1c (BDCA-1) Phycoerythrin (Clone REA694) | Miltenyi | Cat#130-110-536 |
| Anti-human HLA-DR Brilliant Violet 785 (Clone L243) | Biolegend | Cat#307642 |
| LIVE/DEAD Fixable Aqua Dead Cell Stain Kit (405 nm excitation) | Thermo Fisher | Cat#L34957 |
| BUB395 Rat Anti-Integrin B7 | BD Bioscience | Cat#744014 |
| Phosphate Buffered Saline (PBS) | Thermo Fisher | Cat#10010023 |
| Fetal Bovine Serum (FBS) | Wisent | Cat#091-150 |
| Sodium Azide | Bioshop | Cat#SAZ001.250 |
| ALDEFLUOR Kit | Stem Cell | Cat#01700 |
| ALDEFLUOR Assay Buffer | Stem Cell | Cat#01702 |
| Formaldehyde solution 37 wt. % in H2O | Sigma | Cat#F1635-500ML |
| **ELISA** | | |
| Retinoic Acid | MyBioSource | MBS286231 |
| Retinol Binding Protein 4 | R&D System | DRB400 |
| Chemerin | R&D System | DCHM00 |
| I-FABP | Hycult Biotech | HK406 |
| **Software** | | |
| FlowJo version 10 | BD | https://www.flowjo.com/ |
| GraphPad Prism 10 | GraphPad | https://www.graphpad.com/ |
| Cell Engine | CellCarta | https://cellengine.com/#/ |
